# Live-cell Imaging Analysis of Antimycin-Type Depsipeptides via Bioorthogonal Stimulated Raman Scattering Microscopy

**DOI:** 10.1101/583252

**Authors:** Jeremy Seidel, Yupeng Miao, William Porterfield, Wenlong Cai, Xuejun Zhu, Seong-Jong Kim, Fanghao Hu, Santi Bhattarai-Kline, Wei Min, Wenjun Zhang

## Abstract

Small-molecule natural products have been an essential source of pharmaceuticals to treat human diseases, but very little is known about their behavior inside dynamic, living human cells. Here, we demonstrate the first structure-activity-distribution study of complex natural products, the anti-cancer antimycin-type depsipeptides, using the emerging bioorthogonal Stimulated Raman Scattering (SRS) Microscopy. Our results show that the intracellular enrichment and distribution of these compounds are driven by their potency and specific protein targets, as well as the lipophilic nature of compounds.

---

Nature’s small molecules have played an enormous role in the history of medicinal and pharmaceutical chemistry. For example, it has been estimated that over 70% of anti-cancer small molecule treatments are natural products, their derivatives or mimics.^[1]^ Although the pharmaceutical value of natural products has been widely recognized, it is still difficult to transform medicinally active natural products into drugs. One of the major challenges is to understand the complex interplay between natural products and the network of cellular machinery beyond the specific protein targets.^[2]^ This has spurred the development of advanced imaging techniques to obtain views of natural products in cells, but often in a static and destructive manner using bulky fluorescent probes.^[3]^ An improved imaging technique, which provides dynamic views of natural product uptake and distribution in live cells, will have a profound impact on natural product-based drug discovery and development.^[4]^ Tagging natural products with a bio-orthogonal alkyne functionality coupled with the Stimulated Raman Scattering (SRS) microscopy offers such promise.^[5]^

Raman imaging has evolved greatly over the past decade, with improved sensitivity, resolution, and scanning speeds offered by the latest SRS technology (Figure 1).^[6]^ More importantly, compared with other vibrational imaging platforms such as Coherent Anti-Stokes Raman Scattering microscopy, SRS is free of spectral distortion and non-resonant background thus enables quantitative determination of Raman reporters. The vibrational Raman reporter can be as simple as an alkyne,^[5a, 7]^ delivering chemical specificity and biocompatibility for natural product visualization and quantification in complex living systems with minimal activity perturbation of compounds. Compared to fluorescent imaging, SRS imaging offers additional advantages of minimal phototoxicity and photobleaching, allowing prolonged dynamic imaging of tagged natural products within live cells. The SRS microscopy has recently been used to image various alkyne-tagged small-molecule derivatives that have high local intracellular concentrations.^[8]^ Despite the great potential, few natural products have been imaged using SRS microscopy to probe their intracellular behavior.

**Figure 1.**
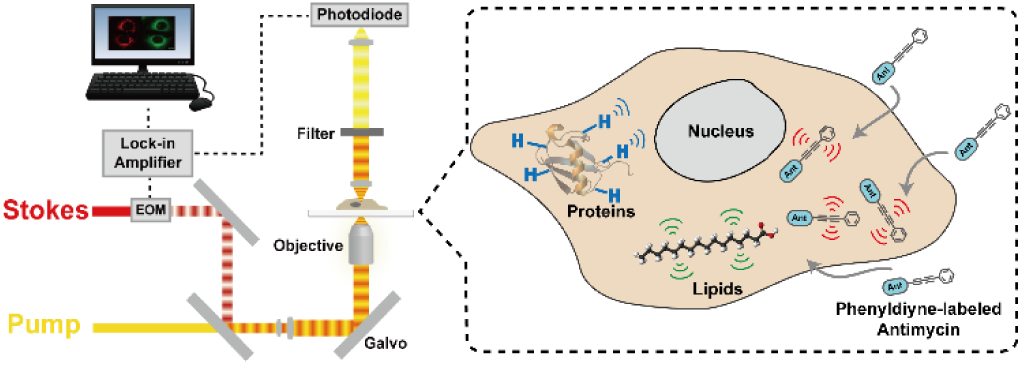
Schematic representation of Stimulated Raman Scattering (SRS) microscopy setup. The Stokes and pump beams are temporally and spatially sychronized while the Stokes beam is modulated with a frequency of 8 MHz. The beams are guided onto the sample by a laser scanning microscope. The transmitted and detected pump lose signal is demodulated by a lock-in amplifier.

Here, we’ve applied SRS imaging to study antimycin-type depsipeptides, a class of complex natural products that have attracted recent attention due to their anti-cancer potential.^[9]^ This family of natural products share a common structural skeleton consisting of a macrocyclic ring with an amide linkage to a 3-formamidosalicylate unit, and primarily differ in the size of their macrolactone ring (Figure 2a). The well-recognized members of this family are the 9-membered antimycins, for which multiple modes of action have been proposed, including inhibition of mitochondrial electron transport chain,^[10]^ anti-apoptotic proteins Bcl_2_/Bcl-x_L_,[11] K-Ras plasma membrane localization,[12] and ATP citrate lyase activity.^[13]^ The levels of contributions of these different mechanisms are unclear. Much less is known about the 15-membered neoantimycins, despite the fact that they have also shown promising anti-cancer activities toward various cancer cell lines.^[14]^ The inhibitory activity of K-Ras plasma membrane localization was shown to be shared between antimycins and neoantimycins,[12] but neoantimycins lacked the Bcl-xL inhibitory activity and were demonstrated to inhibit the expression of GRP-78,^[15]^ a molecular chaperone in the endoplasmic reticulum (ER) that promotes protein folding and provides resistance to both chemotherapy and hypoglycemic stress.^[16]^ To gain additional insights into anti-cancer activities of antimycin-type depsipeptides, we performed a structure-activity-distribution study of both antimycin and neoantimycin against live cancer cells.

**Figure 2.**
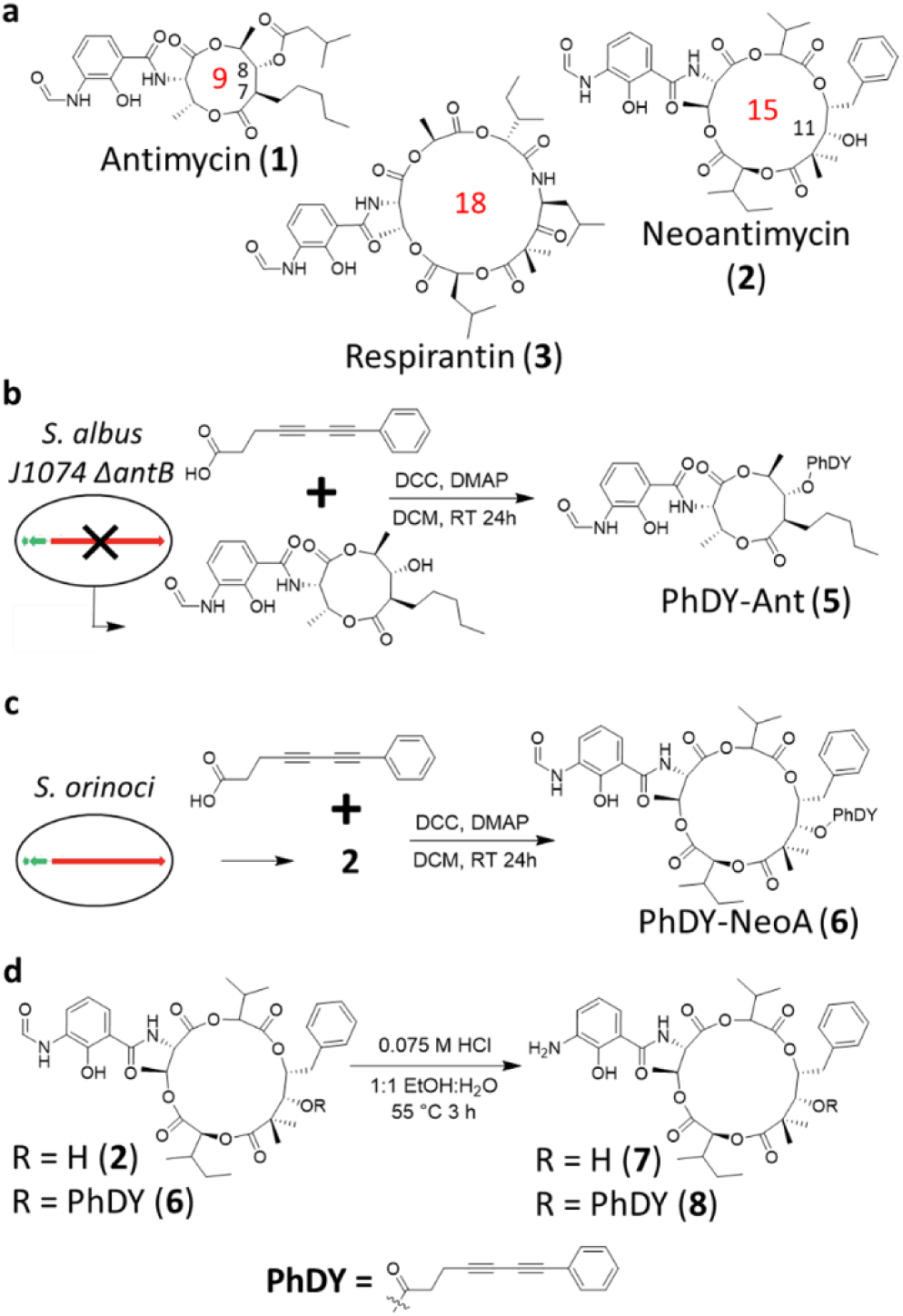
Antimycin-type depsipeptides and their alkyne-tagged derivatives. a) Selected examples of natural antimycin-type depsipeptides. The ring size is indicated in red numbers. b) Generation of 5 through both bioengineering and chemical derivatization. c) Generation of 6 through C-11 esterification. d) Generation of 7 and 8 through acid degradation of 2 and 6, respectively.

To increase the sensitivity of SRS imaging toward bioactive small molecules which are typically used in the low to mid micromolar range,^[17]^ we chose a conjugated diyne with a terminal phenyl ring as a Raman tag. This tag is known to possess increased Raman scattering cross section due to conjugation within the poly-yne chain and the presence of an aryl end-capping group also improves the stability of poly-ynes.^[8]^ This tag has recently been used in separate studies to image a phenyl-diyne anisomycin derivative in mammalian cells and to track the distribution of a cholesterol derivative in *Caenorhabditis elegans* where a detection limit of ∼30 μM was attained.^[8, 18]^ Since bioactive antimycins naturally have high structural variations at the C-7 alkyl and C-8 acyloxy moieties (Figure 2a) and the previous introduction of an alkyne side chain at C-8 did not significantly change cytotoxic activity nor binding of compounds to cancer cells,^[3]^ we reasoned that the Raman tag could be readily introduced at this position with minimal functional perturbation. To prepare phenyl-diyne antimycin (PhDY-Ant, **5**), C-8 deacylated antimycin (**4**) was first purified from the culture of *Streptomyces albus* Δ*antB* in which the last step of C-8 acyloxy formation is abolished in antimycin biosynthesis due to the deletion of the dedicated C-8 acyltransferase AntB.^[19]^ A phenyl-diyne carboxylic acid was chemically synthesized and then coupled to the purified **4** via Steglich esterification to yield **5** (Figure 2b **and S1**). As expected, the MTT proliferation assays with both HeLa (human cervical cancer) and MCF-7 (human breast cancer) cell lines confirmed that PhDY-Ant retained a comparable activity to the natural antimycin (Table 1, **Figure S2**).

**Table 1.** GR50 values (μM) for selected antimycins and neoantimycins against two cancer cell lines in vitro.

| Compound | HeLa | MCF-7 |
| --- | --- | --- |
| Antimycin (**1**) | $20.2 \pm 4.3^a$ | $2.1 \pm 0.5$ |
| PhDY-Ant (**5**) | $31.8 \pm 10.4$ | $28.8 \pm 9.7$ |
| Neoantimycin (**2**) | $38.7 \pm 8.2$ | $24.4 \pm 5.5$ |
| PhDY-NeoA (**6**) | $> 100^b$ | $40.9 \pm 22.8$ |
| Deformylated neoantimycin (**7**) | $> 1000$ | $> 1000$ |

[a] All data were collected in at least triplicate and GR50 values were averaged between at least three biological replicates with error given in standard error of the mean (SEM).
[b] GR50 could not be determined due to solubility limitations.

PhDY-Ant (**5**) was then incubated with HeLa cells and SRS images were acquired by tuning the frequency difference between the pump and Stokes lasers to be resonant with intracellular components such as proteins (CH_3_, 2940 cm^−1^) and lipids (CH2, 2845 cm^−1^), and in the bio-orthogonal region of the Raman spectrum (alkyne, 2251 cm^−1^; off-resonance, 2000 cm^−1^) (Figure 1). **5** was used at solution concentrations ranging from 1-100 µM to probe the detection limit. Intracellular signal could be distinguished at concentrations as low as 10 µM, and contrast was dramatically improved by increasing the solution concentration of **5** to 50 µM (Figure 3 **and S3**). The absolute intracellular concentration of **5** was determined to be ∼ 1.74 mM from the dosing concentration of 50 µM, showing a 35-fold enrichment of this compound in cells. To confirm that the observed signal was driven by the activity of antimycin, a control experiment was performed by incubating 50 µM PhDY tag with cells. No signal of compound was detected inside cells (**Figure S3**), suggesting that the observed Raman signal was not an off-target affect caused by the PhDY tag alone. We next probed the antimycin uptake rate and mechanism. Time-resolved imaging of **5** uptake into live HeLa cells showed that compound uptake was nearly immediate, reaching 75% of the maximum within six minutes (**Figure S4**). **5** appeared to rapidly distribute throughout the cytoplasm of the cells and persist through prolonged incubation. In addition, a low-temperature (4°C) uptake study was performed to investigate possible mechanisms of compound uptake. **5** was absorbed at comparable levels at both 4°C and 37°C (**Figure S5**), suggesting that PhDY-Ant may cross the cell membrane through passive diffusion.

**Figure 3.**
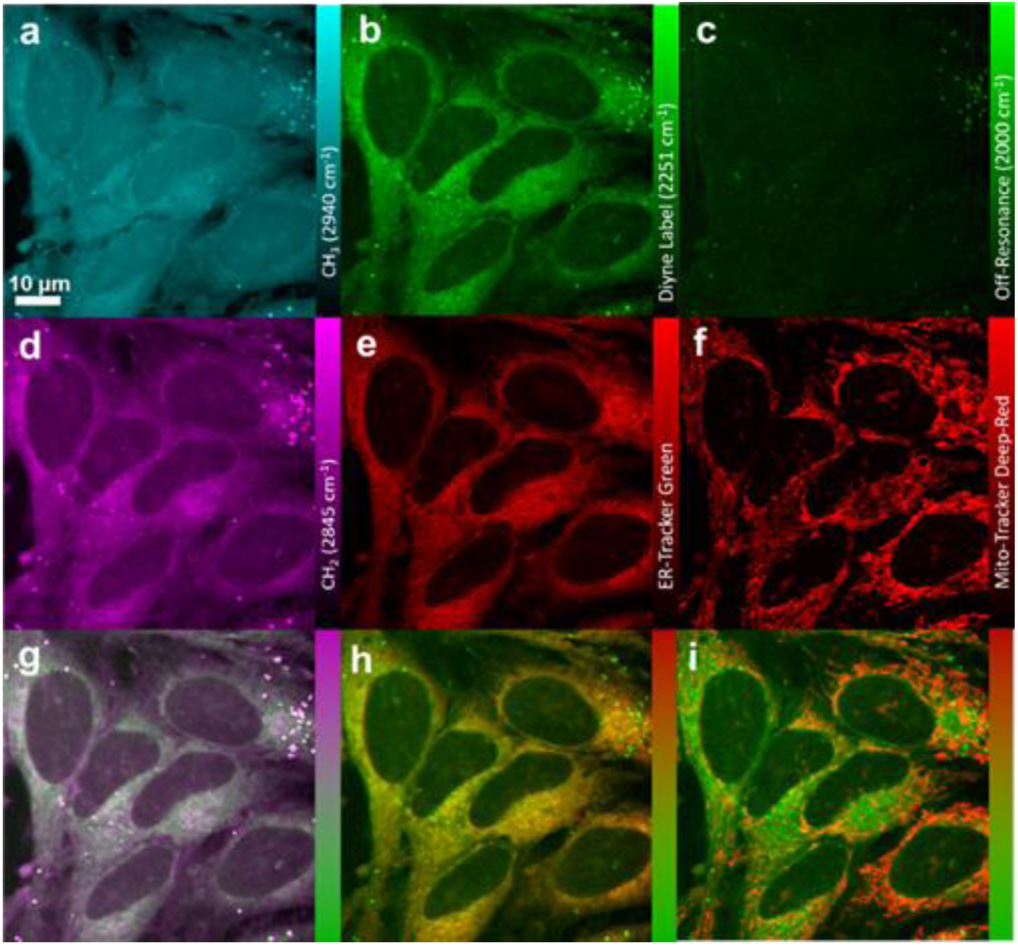
SRS and fluorescence imaging of PhDY-Ant (5) in HeLa cells. a) CH_3_ channel at 2940 cm^−1^ representing proteins. b) Diyne label at 2251 cm^−1^. c) Off-resonance channel at 2000 cm^−1^. d) CH_2_ channel at 2845 cm^−1^, representing lipids. e) Confocal fluorescence imaging of ER-Tracker excited at 488 nm. f) Confocal fluorescence imaging of Mito-Tracker excited at 635 nm. g) Overlay image of d) lipids and b) diyne label. h) Overlay image of e) ER-Tracker and b) diyne label. i) Overlay image of f) Mito-Tracker and b) diyne label.

The non-destructive nature of SRS imaging allows follow-up studies on compound distribution inside the cell, as well as correlating compound uptake with any possible phenotypic changes to cellular composition using dual-color and multi-modal approaches. For example, using a multi-modal approach to probe the subcellular localization of **5**, HeLa cells were treated with ER-Tracker Green and Mito-Tracker Deep Red, cell-permeable fluorescent stains selective for the endoplasmic reticulum and mitochondria, respectively. Inspection of the merged images demonstrated that **5** correlated well with ER-Tracker (Figure 3). This colocalization agrees with one of its known direct protein targets, Bcl_2_, an anti-apoptotic protein localized primarily in the ER.^[20]^ Notably, a prolonged incubation led to a decrease of correlation between ER-Tracker and **5** (**Figure S6**), possibly due to compound dissociation from targets and/or translocation of protein targets. In spite of the known activity of antimycin in inhibiting the mitochondrial electron transport chain by binding to the quinone reduction site Q_i_ of the cytochrome *bc*_1_ complex,[10] no significant colocalization of **5** with Mito-Tracker was found (Figure 3). In addition, no obvious phenotypic changes were observed in lipids or proteins during the eight-hour incubation, although the merged images for **5** and the lipid channel showed a strong correlation, especially in lipid droplets (Figure 3). This correlation is likely caused by the lipophilic nature of **5** rather than any specific binding. Further image analysis using profile plots demonstrated that the localization of **5** was best explained by a combination of ER-Tracker and lipids (**Figure S7**). This result indicates that the intracellular behavior of antimycin is not totally dictated by specific protein binding nor non-specific absorption. Instead, there is likely a complex interplay between antimycin, its protein targets, and the lipid-rich regions of the cell.

We next analyzed the distribution of PhDY-Ant (**5**) in MCF-7 and compared it to HeLa cells to probe if the distribution is cell-line specific. Similar to HeLa cells, **5** colocalized with ER-Tracker, but not Mito-Tracker (**Figure S8**). These data suggest that localization of antimycin in the ER is conserved across different cancer cell lines. In addition, MCF-7 cells showed a much smaller number of lipid droplets than HeLa cells, but instead contained highly lipid-rich regions at the intercellular boundaries that did not attract **5** (**Figure S8**). This result further suggests that the characteristics and location of lipids may also be important for enrichment of antimycin. Indeed, a profile analysis showed that certain areas of localization of **5** in MCF-7 cells were best correlated with ER-Tracker while others were best correlated with the lipid channel (**Figure S9**).

Compared to antimycin, the molecular mechanism for the ring-expanded neoantimycin to inhibit cancer cell growth is much less known, and no direct protein target of neoantimycin has been identified. In addition, limited structure-activity relationship studies have been performed on the molecular scaffold of neoantimycin. It is yet to be determined if a similar tagging strategy, the esterification of the macrolactone C-11 hydroxyl moiety that is naturally present in neoantimycin (Figure 2a), can be adopted to produce a neoantimycin derivative that is suitable for imaging analysis while retaining its anti-cancer activity. Neoantimycin was purified from the culture of *S. orinoci* and subjected to esterification by a phenyl-diyne carboxylic acid to generate phenyl-diyne neoantimycin (PhDY-NeoA, **6**) (Figure 2c and **S10**). The MTT proliferation assays with both HeLa and MCF-7 cells indicated that **6** had a slightly decreased but significant bioactivity, although its GR50 value against HeLa cells could not be determined due to solubility limitation (Table 1, **Figure S11**). This response of cell lines to the C-11 modification is consistent with a recent report in which oxidation of the same hydroxyl to ketone of neoantimycin led to slightly increased IC_50_ values against multiple cancer cell lines.^[14]^ The *N*-formyl group has been conserved in the antimycin-type depsipeptides and linked to respiration inhibition for antimycin.^[9]^ To probe the role of the *N*-formyl group in anti-cancer activities of neoantimycin, we produced deformylated neoantimycin (**7**) and its tagged version (**8**) through acid degradation of **2** and **6**, respectively (Figure 2d, **S12 and S13**). The growth of HeLa and MCF-7 cells was not inhibited upon treatment of up to 1 mM of **7** (Table 1 and **Figure S11**), demonstrating the critical role of this moiety for anti-cancer activity of the 15-membered neoantimycin. This is in contrast to the 18-membered antimycin-type depsipeptides (Figure 2a) of which deformylation did not significantly decrease the inhibitory activity toward various cancer cell lines,^[21]^ suggesting different modes of action for 15-and 18-membered compounds. Nonetheless, the generation of both active and inactive tagged neoantimycins provided an opportunity to investigate possible differential uptake of these compounds in live cells.

PhDY-NeoA (**6**) and deformylated PhDY-NeoA (**8**) were then subjected to SRS imaging analysis with both HeLa and MCF-7 cells. Enrichment of both compounds in lipid droplets was observed for both cell lines, which is likely due to the lipophilic nature of compounds rather than any specific binding to targets (**Figure S14**). In addition, **6** was detected by SRS throughout the cytoplasm of the MCF-7 cells with an estimated intracellular concentration of ∼ 0.82 mM from a dosing concentration of 100 µM. **6** was also detected at similar concentrations throughout the cytoplasm of the HeLa cells, although with a decreased signal intensity, demonstrating a positive correlation of compound intracellular enrichment to its cytotoxic activity. This is particularly true for intracellular enrichment of **8**, for which very weak SRS signals were detected (except within lipid droplets) in either cell line (**Figure S14**). Further analysis using dual-color and multi-modal approaches showed that the intracellular distribution of PhDY-NeoA (**6**) differed from that of PhDY-Ant (**5**). In particular, **6** showed no correlation with Mito-Tracker and a weak correlation with ER-Tracker in MCF-7 cells (Figure 4). Line plot analysis showed that **6** was much better correlated with lipids than with ER-Tracker (**Figure S15**), suggesting that neoantimycin may not target ER. This observation is consistent with previous reports that the known antimycin target, Bcl2/Bcl-xL, is not a neoantimycin protein target.^[22]^

**Figure 4.**
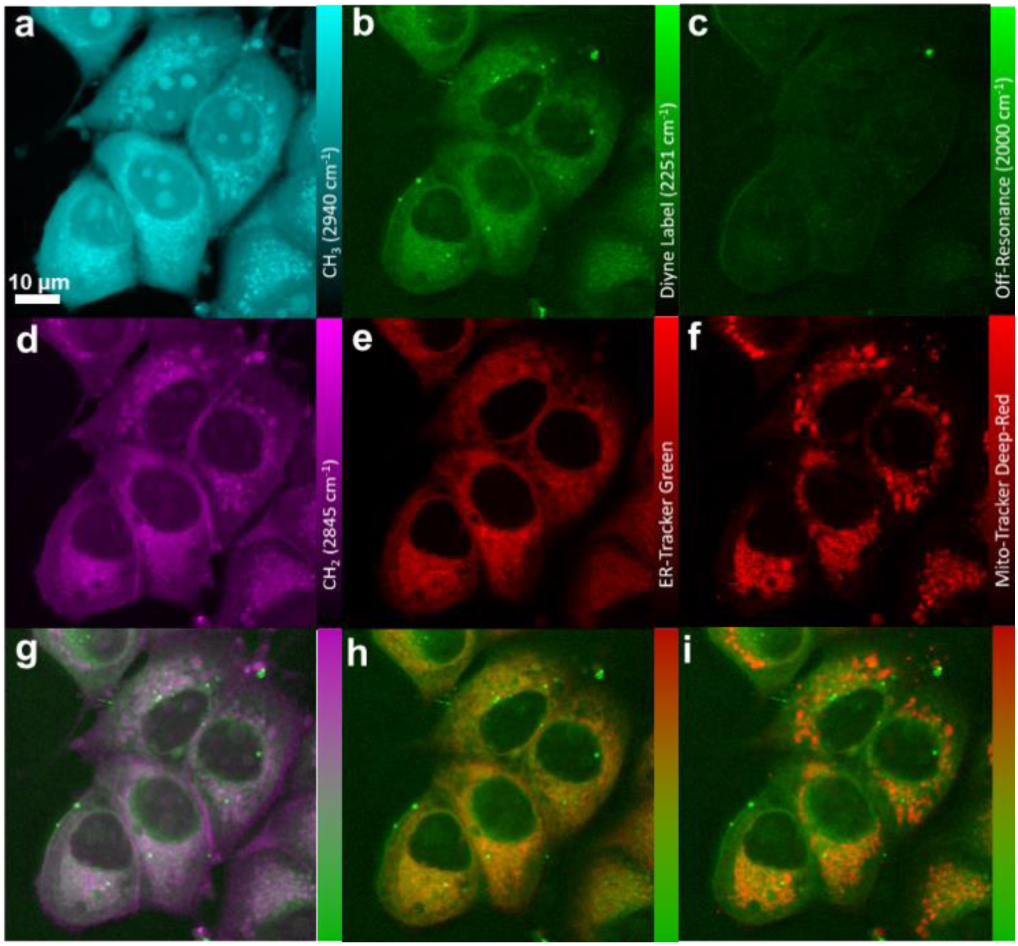
SRS and fluorescence imaging of PhDY-NeoA (6) in MCF-7 cells. a) CH3 channel at 2940 cm^−1^ representing proteins. b) Diyne label at 2251 cm^−1^. c) Off-resonance channel at 2000 cm^−1^. d) CH2 channel at 2845 cm^−1^, representing lipids. e) Confocal fluorescence imaging of ER-Tracker excited at 488 nm. f) Confocal fluorescence imaging of Mito-Tracker excited at 635 nm. g) Overlay image of d) lipids and b) diyne label. h) Overlay image of e) ER-Tracker and b) diyne label. i) Overlay image of f) Mito-Tracker and b) diyne label.

In summary, both the 9-and 15-membered antimycin-type depsipeptides have been labeled with an alkyne Raman tag and subjected to bioorthogonal SRS imaging analysis in live cancer cells. This work provides the first global and dynamic view of the interplay between these anti-cancer complex natural products and the complicated network of cellular machinery. We demonstrate the primary localization of the 9-membered antimycin in the endoplasmic reticulum despite the previous known protein targets of antimycin in various cellular organelles. We further show that the anti-cancer activity of the 15-membered neoantimycin is dependent on the *N*-formyl moiety and less sensitive toward the C-11 modification. A different cellular localization of neoantimycin compared to antimycin is also indicated. Our results suggest that the intracellular enrichment and distribution of these compounds are driven by their potency and specific protein targets, as well as the lipophilic properties of compounds. This non-destructive imaging technique of drug candidates is expected to complement existing biochemical and proteomic techniques in the early stages of drug discovery, facilitating efforts in reducing off-target effects and improving efficacy of candidate compounds.

## Experimental Section

Full experimental details are available in the supporting information.

## Acknowledgements

J.A.S. is supported by the National Science Foundation Graduate Research Fellowship Program. This research was financially supported by grants to W.Z. from the American Cancer Society, Alfred P. Sloan Foundation, and the Chan Zuckerberg Biohub Investigator Program. W. M. acknowledges support of R01EB020892 and R01GM128214 from NIH, and the Camille and Henry Dreyfus Foundation. We thank the Berkeley Cell Culture Facility for providing cell culturing services for the cytotoxicity assays.

