## Supplementary Information for "Live-cell Imaging Analysis of Antimycin-Type Depsipeptides via Bioorthogonal Stimulated Raman Scattering Microscopy"

**Supporting Information**

**Table of Contents**

**Experimental Section**

**Supplementary Figures**

**Figure S1.** NMR, MS, and UV-vis characterization of PhDY-Ant (**5**).

**Figure S2.** PhDY-Ant (**5**) and antimycin (**1**) dose response curves.

**Figure S3.** Intracellular PhDY-Ant (**5**) and phenyl-diyne tag imaging at various concentrations.

**Figure S4.** Time course of PhDY-Ant (**5**) uptake in HeLa cells.

**Figure S5.** Temperature dependent uptake of PhDY-Ant (**5**).

**Figure S6.** Time-dependent colocalization of PhDY-Ant (**5**) in HeLa cells.

**Figure S7.** Line plot analysis of PhDY-Ant (**5**) in HeLa cells.

**Figure S8.** Intracellular imaging of PhDY-Ant (**5**) in MCF-7 cells.

**Figure S9.** Line plot analysis of PhDY-Ant (**5**) in MCF-7 cells.

**Figure S10.** NMR, MS, and UV-vis characterization of PhDY-NeoA (**6**).

**Figure S11.** Dose response curve for various neoantimycins in MCF-7 cells.

**Figure S12.** NMR, MS, and UV-vis characterization of deformylated neoantimycin (**7**).

**Figure S13.** NMR, MS and UV-vis characterization of deformylated PhDY-NeoA (**8**).

**Figure S14.** Imaging of differential uptake of PhDY-NeoA (**6**) and deformylated PhDY-NeoA (**8**) in HeLa and MCF-7 cells.

**Figure S15.** Line plot analysis of PhDY-NeoA (**6**) in MCF-7 cells.

### Experimental Section

#### Spontaneous Raman Spectroscopy

All spontaneous Raman spectra were acquired with a commercial Raman microscope (Xplora, Horiba Jobin Yvon). All diyne labelled compounds were dissolved in dimethyl sulfoxide (DMSO) with a concentration of 50 mM for Raman spectrum acquisition. A 532 nm diode laser beam was guided through a 50×, 0.75 N.A. air objective (NPLAN EPI, Leica) to excite samples. All spectra were obtained with 5 s acquisition time and 10 times averaging. All Raman spectrum data were processed with LabSpec 6 software.

#### General cell culture

MCF7 and HeLa cells were obtained from the UCB Cell Culture Facility cultured in DMEM (Gibco) supplemented with 10% (vol/vol) fetal bovine serum, GlutaMAX, penicillin, and streptomycin. Cells were maintained at 37 °C with 5% $CO_2$ in a water saturated incubator.

#### Sample preparation for SRS and fluorescence imaging of live cells

HeLa and MCF-7 cells were seeded on round coverslips with 800 µL complete growth media in four-well plates 2 days before the experiment. On the day of the experiment, the culture media was replaced with 500 µL fresh culture media supplemented with fluorescent organelle markers (MitoTracker™ Deep Red FM, Invitrogen; ER-Tracker™ Green (BODIPY™ FL Glibenclamide), Invitrogen) and diyne-labelled compounds at specific concentrations. The cells were incubated for designated time in the incubator until washed with phosphate buffered saline (PBS, Gibco) three times. The coverslips with cells were placed onto imaging spacers (SecureSeal™ imaging spacers, Sigma-Aldrich) and attached to glass slides. The space between coverslips and glass slides was filled with PBS solution to preserve the live cells for a short period.

#### Confocal Fluorescence Microscopy

All fluorescence imaging was performed with a commercial Olympus FV1200 confocal microscope with standard laser excitation and bandpass filter set for ER-Tracker and Mito-Tracker. The objectives were either a 25× 1.05 N.A. water objective (XLPlan N, Olympus) or a 60× 1.2 N.A. water objective (UPlanAPO/IR, Olympus). The images were acquired with Olympus FV10 software and processed with ImageJ.

#### Stimulated Raman Scattering (SRS) Microscopy

SRS imaging was performed with the same microscope setup as the fluorescence imaging but with customized laser input. A commercial laser source (picoEmerald, Applied Physics & Electronics, Inc.) was utilized to produce both the Pump and Stokes beams for SRS. The wavelength of Stokes beam was 1064 nm. The beam was modulated at 8 MHz by an electro-optic modulator (EOM). The Pump beam had tunable wavelengths within a range of 720 nm to 990 nm. Both beams had 6 ps pulse width and 80 MHz repetition rate. The two beams were spatially and temporally overlapped and tightly focused onto the sample, after which Pump loss and Stokes Gain were generated. Both transmitted beams were collected by a 1.4 N.A. oil condenser. The Stokes beam was filtered off with a high O.D. bandpass filter (890/220 CARS, Chroma Technology), while the Pump beam was detected by a silicon photodiode (FDS1010, Thorlabs) with a DC voltage of 64 V. The output current was terminated by a 50 Ω terminator and demodulated by a lock-in amplifier (SR844, Stanford Research Systems) at 8 MHz frequency.

The Pump loss signal at each pixel was sent to the FV10 analog channel to generate the images. To obtain sufficient resolution and sensitivity, all live cell images were acquired with 100 us time constant and 512 × 512 pixel number. The $CH_3$ channel at 2940 $cm^{-1}$ was obtained with 40 mW Pump power and 100 mW Stokes power on sample. The $CH_2$ channel at 2845 $cm^{-1}$, diyne channel at 2251 $cm^{-1}$ and off-resonance channel at 2000 $cm^{-1}$ were all obtained with 100 mW Pump power and 100 mW Stokes power on sample.

**Image Analysis**

All images were generated by Olympus FV 10 software. Images were taken under strictly the same conditions (pump and Stokes power, digital gain and offset) when comparison is performed.

The Raman beams and the fluorescent beams are not in-line in the microscope set-up. As a result, the two modalities have a slight offset in the images that they acquire. Before any image analysis or background reduction was performed, this misalignment had to be corrected. MATLAB code was used to align the images to achieve the highest possible colocalization coefficient between the Raman and fluorescent channels. The non-overlapping regions of the different image channels were then cropped off immediately. Subsequent analysis of overlaid images showed that the many channels were very well-aligned.

All image analysis was performed in FIJI. Because the images had varying levels of background, some background reduction had to be performed to facilitate the analysis. Every image channel (except for -CH3) of every image set was subjected to an identical rolling ball background reduction with a radius of 200 pixels. This relatively large radius ensured against dimming or outright removing features with a homogenous intensity. The relative brightness of images were occasionally adjusted via the multiply function in imageJ prior to merging the images. This was purely to help the reader observe colocalization between two channels at different intensities. The image intensities were not modified for any non-merged images.

**Cell viability assays**

HeLa or MCF7 were grown in clear, flat bottom 96 well plates (Olympus plastics). MTT cell growth assay reagents (EMD Millipore) were used as purchased. In brief, to cells in 100 µL DMEM media, 10 µL of MTT solution was added. The assay was allowed to develop at 37 °C for 4 h. After MTT formazan formation 100 µL 0.04 M HCl in isopropanol was added to each well and mixed until all solid MTT formazan was dissolved. The 96-well plates were read for absorbance at 570 nm using a Tecan M1000 plate reader.

For cytotoxicity studies, compounds, dissolved in DMSO, were added to wells (final DMSO concentration of 0.5-1%, referenced to control wells containing DMSO alone) at 20% cell confluency. Cells were incubated with compounds for 48 h at 37 °C prior to MTT addition.

Growth response (GR) values were obtained by getting a baseline cell count using the MTT assay performed concurrently with cytotoxic compound addition in a separate plate as described by Hafner *et al.*[1] GR50 values and plots were obtained using the GR metrics software package accessed through github (https://github.com/sorgerlab/gr50_tools).[1] All data were collected in at least triplicate and GR50 values were averaged between at least three biological replicates with error given in standard error of the mean (SEM).

**General synthetic methods**

All reagents were purchased from commercial suppliers and used without further purification. Reaction progress was monitored by thin-layer chromatography on silica gel 60 plates (aluminum back, EMD Millipore) and visualized by UV light or by LC/MS (Agilent 6120 single quadrupole). Samples were analyzed by reverse phase HPLC on a C18 column (Eclipse Plus 3.5 µM 100 x 4.6 mm) with the following elution protocol: A 40-80% MeCN in $H_2O$ (0.1% FA) linear gradient over 3 min, followed by a 80-100% MeCN in $H_2O$ (0.1% FA) over 22 min with a flow rate of 0.5 mL/min. Compounds were purified by flash column chromatography using Fisher Scientific 230-400 mesh, 60 Å, silica gel. HPLC purifications were performed on an Agilent 1260 HPLC with a semipreparative scale were performed using an Altima C18 column 5 µM (150 x 10 mm). NMR spectra were acquired with a Bruker Biospin spectrometer with a cryoprobe. All spectra were acquired at 298 K. $^1$H spectra were acquired at 900 MHz, $^{13}$C spectra were acquired at 226 MHz. Coupling constants (*J*) are provided in Hz and chemical shifts reported in ppm relative to residual non-deuterated NMR solvent. High resolution mass spectra were collected using an Agilent Technologies 6520 Accurate-Mass Q-TOF LC-MS instrument. UV data was obtained on an Agilent 1260 series DAD.

**Phenyl-diyne acid synthesis**

The phenyl-diyne acid tag was synthesized according to the method described by Balaraman *et. al*.[2] First, $Cu(Ac)_2$ monohydrate was added to 50 mL of methylene chloride in a round bottom flask until saturation was achieved. Then, 5 equivalents of piperidine was added, followed by 5 equivalents of phenylacetylene and 1 equivalent of 4-pentynoic acid. The mixture was left exposed to air for the duration of the reaction. After the reaction mixture had stirred at room temperature for 8 hours, the excess $Cu(Ac)_2$ was removed by centrifugation. The clarified reaction mixture was then concentrated to 10 mL in vacuo. The $Cu(Ac)_2$ that had precipitated was once again removed by centrifugation. The concentrated, clarified reaction mixture was then fractionated via silica gel chromatography. The fractions containing the acid product were identified via TLC analysis and combined. After evaporating the combined fractions in vacuo, the purity of the product was confirmed via LC-MS and UV-vis analysis.

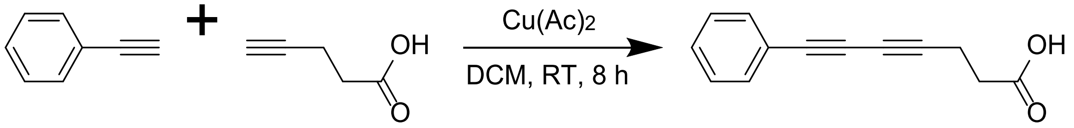

**Deacylated antimycin production**

Deacylated antimycin was produced using an engineered *Streptomyces albus* J1074 strain with the *antB* gene deleted in-frame through double crossover. A seed culture was grown in two separate aliquots of 5 mL tryptic soy broth at 30˚C for 48 hours. Next, 1 mL of the seed culture was added to 10 separate 1 L aliquots of mannitol soy broth in 4 L plastic flasks. After incubating for 6 days at 30˚C and 200 RPM, the solids were removed from the fermentation media via centrifugation. The clarified media was then extracted with 50 g/L of XAD-16 resin. The resin was then washed with 10 L of distilled water and subsequently extracted with 3 L of methanol. The methanol extract was then dried in vacuo and purified by silica gel column chromatography using a gradient of 0% to 100% ethyl acetate in methylene chloride in 25% increments. The fractions containing deacylated antimycin were combined, evaporated, dissolved in methanol and further purified by HPLC on a semi-preparative C18 column (10 mm i.d., 250 mm length, Vydac) with a linear gradient of 25 to 95% acetonitrile (vol/vol) over 20 min, and 95% acetonitrile (vol/vol) for a further 10 min in H2O with 0.1% (vol/vol) formic acid (FA) at a flow rate of 2 mL/minFractions were collected manually and concentrated under vacuum. Fractions containing *S. albus* Δ*antB* antimycins were pooled and further separated by HPLC using an Agilent Eclipse Plus C18 column (4.6 x 100 mm) with a linear gradient of 25 to 95% acetonitrile (vol/vol) over 20 min, and held at 95% acetonitrile (vol/vol) for 10 min in H2O with 0.1% (vol/vol) formic acid (FA) at a flow rate of 1 mL/min. Total yield was roughly 5 mg/L.

**Neoantimycin production**

Neoantimycin was produced using a wild-type strain of *Streptomyces orinoci*. *S. orinoci* was grown three separate 5 mL cultures of TSB for 72 hours. After that 50 µL of seed culture was added to 40 separate 150 mL plastic shake flasks each containing 50 mL of mannitol soy broth. After 6 days of fermentation, the broth was clarified by centrifugation. The 2 L of broth was extracted two times consecutively with 2 L of ethyl acetate. The ethyl acetate extract was dried with sodium sulfate and evaporated in vacuo. After redissolving the residue in 20 mL of ethyl acetate, it was fractionated by silica gel chromatography using a gradient of 0% to 100% ethyl acetate in methylene chloride in 25% increments. The fractions with neoantimycin were identified via LC-MS and combined. The fractions containing neoantimycin were combined, evaporated, dissolved in methanol and further purified by HPLC on a semi-preparative C18 column (10 mm i.d., 250 mm length, Vydac) with a linear gradient of 25 to 95% acetonitrile (vol/vol) over 20 min, and 95% acetonitrile (vol/vol) for a further 10 min in $H_2O$ with 0.1% (vol/vol) formic acid (FA) at a flow rate of 2 mL/min. Fractions containing *S. orinoci* neoantimycins were pooled and further separated by HPLC using an Agilent Eclipse Plus C18 column (4.6 x 100 mm) with a linear gradient of 25 to 95% acetonitrile (vol/vol) over 20 min, and held at 95% acetonitrile (vol/vol) for 10 min in $H_2O$ with 0.1% (vol/vol) formic acid (FA) at a flow rate of 1 mL/min. Total yield was roughly 10 mg/L.

**Esterification**

Deacylated antimycin (4.00 mg and 0.00886 mmol) and neoantimycin (6.20 mg and 0.00886 mmol) were converted to PhDY-Ant and PhDY-NeoA respectively via Steglich esterification with phenyl-diyne acid. Phenyl-diyne acid (1.93 mg and 0.00975 mmol), DCC (2.01 mg and 0.00975 mmol), and DMAP (1.19 mg and 0.00975 mmol) were added to methylene chloride (5 mL) stirred in a round bottom flask and chilled in an ice bath. The round bottom was then evacuated and flushed with and kept under inert atmosphere. After stirring for one hour, the alcohol (either deacylated antimycin or neoantimycin) was added to the flask. The reaction was allowed warm to room temperature and stirred for 24 hours. The reaction mixture was then extracted three times with water. The fractions were combined and back-extracted with methylene chloride. The two methylene chloride fractions were then combined and evaporated in vacuo. After dissolving the product in methanol, it was purified via HPLC using an Agilent Eclipse Plus C18 column (4.6 x 100 mm) with a linear gradient of 55 to 95% acetonitrile (vol/vol) over 20 min, and 95% acetonitrile (vol/vol) for a further 10 min in $H_2O$ with 0.1% (vol/vol) formic acid (FA) at a flow rate of 2 mL/min. Total yield was roughly 70% for PhDY-Ant and 35% for PhDY-NeoA.

**PhDY-Ant (5)**

$^1$H NMR (900 MHz, DMSO-$d_6$) $\partial$ 12.50 (s, 1H), 9.59 (s, 1H), 9.26 (d, $J$ = 8.3 Hz, 1H), 7.96 (d, $J$ = 7.9 Hz, 1H), 7.88 (d, $J$ = 6.6 Hz, 1H), 7.53 (d, $J$ = 7.2 Hz, 2H), 7.44 (t, $J$ = 7.5 Hz, 1H), 7.40 (t, $J$ = 7.7 Hz, 2H), 6.93 (t, $J$ = 7.9 Hz, 1H), 5.68 (d, $J$ = 8.4 Hz, 1H), 5.52 (p, $J$ = 6.8 Hz, 1H), 5.25 (app t, $J$ = 7.9 Hz, 1H), 4.67 (dq, $J$ = 9.5, 6.3 Hz, 1H), 3.31-3.30 (overlap m, $J$ = 8.6 Hz, 1H), 2.72 (d, $J$

= 3.8 Hz, 4H), 2.17 (dt, *J* = 10.5, 3.0 Hz, 1H), 2.03 – 1.95 (m, 1H), 1.83 (tdd, *J* = 11.2, 6.2, 2.9 Hz, 1H), 1.51-1.48 (m, 1H), 1.32 (d, *J* = 6.3 Hz, 3H), 1.27 (d, *J* = 6.8 Hz, 3H), 1.09 – 1.02 (m, 2H), 0.85 (d, *J* = 6.6 Hz, 3H), 0.84 (d, *J* = 6.7 Hz, 3H). $^{13}C$ NMR (226 MHz, DMSO-$d_6$) ∂ 174.9, 170.0, 169.8, 168.8, 151.1, 132.4, 129.6, 128.8, 127.2, 127.0, 124.3, 120.7, 118.4, 115.4, 84.9, 76.2, 76.0, 75.0, 74.2, 70.8, 64.9, 53.8, 52.1, 36.1, 34.1, 27.5, 26.5, 22.6, 22.1, 18.4, 15.20, 15.06, 13.97. MS (ESI) calcd for $C_{35}H_{39}N_2O_9$ $[M+H]^+$ *m/z* 631.2650 found 631.2649.

**PhDY-NeoA (6)**

$^1H$ NMR (900 MHz, DMSO-$d_6$) ∂ 12.62 (s, 1H), 9.64 (s, 1H), 9.31 (br s, 1H), 8.13 (s, 1H), 7.98 (d, *J* = 7.9 Hz, 1H), 7.74 (d, *J* = 7.7 Hz, 1H), 7.52 (d, *J* = 7.7 Hz, 2H), 7.44 (t, *J* = 7.3 Hz, 1H), 7.39 (t, *J* = 7.6 Hz, 2H), 7.32 – 7.27 (m, 4H), 7.22 (t, *J* = 6.9 Hz, 1H), 6.92 (br s, 1H), 5.39 – 5.34 (m, 2H), 5.18 (d, *J* = 6.1 Hz, 1H), 5.13 (d, *J* = 3.4 Hz, 1H), 5.08 (dt, *J* = 10.0, 3.6 Hz, 1H), 4.95 (dd, *J* = 8.4, 5.8 Hz, 1H), 3.82 (app t, *J* = 5.4 Hz, 1H), 2.91 (dd, *J* = 14.6, 9.7 Hz, 1H), 2.81 (dd, *J* = 14.5, 3.6 Hz, 1H), 2.73-2.71 (m, 4H), 2.26-2.22 (m, 1H), 1.69-1.66 (m, 1H), 1.36-1.32 (m, 1H), 1.32 (d, *J* = 6.3 Hz, 3H), 1.28 (s, 3H), 1.11 – 1.07 (m, 1H), 1.06 (s, 3H), 1.00 (d, *J* = 6.8 Hz, 3H), 0.88 (d, *J* = 6.9 Hz, 3H), 0.77 (d, *J* = 6.9 Hz, 3H), 0.75 (t, *J* = 7.4 Hz, 2H). $^{13}C$ NMR (226 MHz, DMSO) ∂ 186.5, 184.2, 179.4, 179.3, 173.0, 169.7, 168.4, 163.5, 159.0, 137.5, 135.2, 132.8, 132.6, 130.0, 129.6, 129.3, 128.82, 128.78, 127.0, 121.1, 118.5, 115.6, 95.2, 85.3, 80.0, 79.1, 77.1, 75.4, 74.7, 74.6, 69.5, 62.6, 56.1, 44.6, 38.6, 34.9, 34.5, 30.0, 24.1, 22.5, 19.1, 18.4, 17.3, 16.6, 15.6, 15.5, 11.8. MS (ESI) calcd for $C_{49}H_{55}N_2O_{13}$ $[M+H]^+$ *m/z* 879.3699, found 879.3706.

**Deformylated NeoA (7)**

Neoantimycin (**2**), 0.050 g, was dissolved in 0.25 mL EtOH followed by 0.25 mL 0.03 M HCl and the reaction was stirred at 55 °C for 3 h. The reaction was quenched with 0.80 mL 100 mM ammonium bicarbonate (pH 7) and the EtOH was evaporated *in vacuo*. The solution was then purified by HPLC (semi-preparative, reversed phase using the following elution protocol: 30% MeCN for 2 min, followed by a linear gradient of 30-90% MeCN in $H_2O$ over 24 min, with a flow rate of 1 mL/min) to yield a white solid (0.011 g, 24%). $^1H$ NMR (900 MHz, $CD_3OD$) ∂ 7.3 (app d, J = 4.4, 4H), 7.2 (m, 2H), 6.93 (dd, J = 7.7, 1.5, 1H), 6.72 (t , J = 7.9, 1H), 5.64 (m, 1H), 5.42 (d , J = 3.7, 1H), 5.19 (d , J = 4.2, 1H), 5.1 (d , J = 3.3, 1H), 5 (dt , J = 8.4, 3.9, 1H), 4.04 (d , J = 5.1, 1H), 3.05 (dd , J = 14.7, 9.3, 1H), 2.93 (dd , J = 14.6, 4.1, 1H), 2.34 (m, 1H), 1.8 (m, 1H), 1.5 (m, 1H), 1.4 (d , J = 6.4, 3H), 1.35 (s, 3H), 1.23 (m, 1H), 1.18 (s, 3H), 1.11 (d , J = 6.9, 3H), 0.98 (d , J = 6.8, 3H), 0.92 (d , J = 6.83, 3H), 0.88 (t , J = 7.5, 3H). $^{13}C$ NMR (226 MHz, $CD_3OD$) ∂ 181.4, 174.0, 172.8, 170.6, 169.7, 138.2, 138.1, 130.7, 130.31, 130.30, 129.6, 127.8, 125.2, 124.8, 120.1, 119.7, 81.8, 80.4, 78.6, 76.5, 71.7, 56.8, 46.1, 40.1, 36.3, 31.4, 30.8, 25.2, 23.0, 19.5, 18.9, 17.3, 17.0, 15.6, 12.0. MS (ESI) calcd for $C_{35}H_{47}N_2O_{11}$ $[M+H]^+$ *m/z* 671.3174, found 671.3236.

**Deformylated PhDY-NeoA (8)**

PhDY-NeoA (**7**), 0.0085 g, was dissolved in 0.30 mL EtOH followed by 0.30 mL 0.03 M HCl and the reaction was stirred at 55 °C for 8 h. The reaction was quenched with 0.80 mL 100 mM ammonium bicarbonate (pH 7) and the EtOH was evaporated *in vacuo*. The solution was then purified by HPLC (semi-preparative, reversed phase, using the following elution protocol: 40% MeCN for 2 min, followed by a linear gradient of 40-90% MeCN in $H_2O$ over 24 min, with a flow rate of 1 mL/min) to yield a light brown solid (0.0029 g, 35%). See NMR table below for spectral information. MS (ESI) calcd for $C_{48}H_{55}N_2O_{12}$ $[M+H]^+$ *m/z* 851.3750, found 851.3754.

Deformylated PhDY-NeoA (**8**) NMR table includes 2D NMR data to ensure PhDY tag had not migrated during acid degradation. Atom numbers kept the same as Takeda *et al.*[3] to avoid confusion.

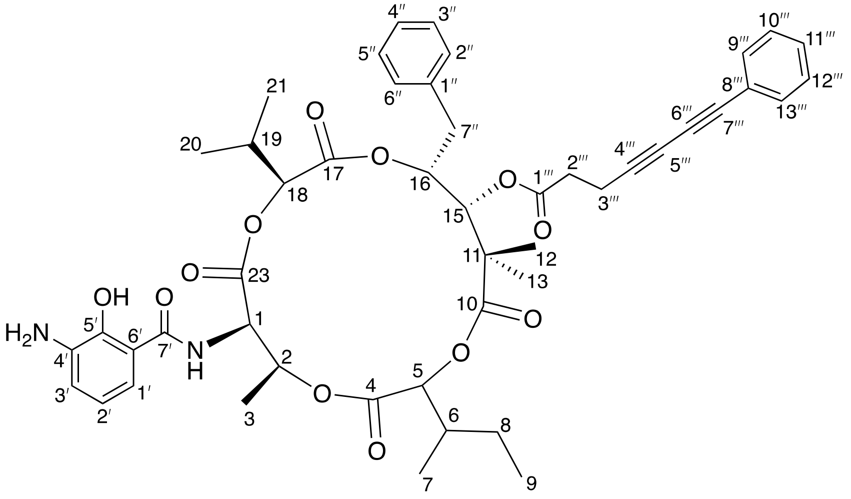

| position | $\delta_H$ [ppm] | integration | mult. ($J$ in Hz) | $^1$H-$^{13}$C HSQC | $^1$H-$^{13}$C HMBC | $\delta_C$ [ppm] |
|---|---|---|---|---|---|---|
| 1 | 5.14 | 1 | br s | 1 | | 57.0 |
| 2 | 5.63 | 1 | m | 2 | | 72.0 |
| 3 | 1.43 | 3 | d (6.32) | 3 | 2, 1, 19, 13 | 17.3 |
| 4 | | | | | | 169.9 |
| 5 | 5.11 | 1 | d (3.3) | 5 | 4, 19, 3 | 78.5 |
| 6 | 1.5 | 1 | m | 8 | 3 | 23.4 |
| 7 | 0.96 | 3 | d (6.86) | 7 | 5, 8 | 15.9 |
| 8 | 1.26 | 1 | m | 8 | | 25.1 |
| | 0.94 | 1 | m | 8 | 7 | |
| 9 | 0.91 | 3 | t (7.46) | 9 | 8 | 12.0 |
| 10 | | | | | | 181.5 |
| 11 | | | | | | 46.0 |
| 12 | 1.18 | 3 | | 12 | 10, 18, 11, 13 | 18.9 |
| 13 | 1.35 | 3 | | 13 | 10, 18, 11, 12 | 23.0 |
| 15 | 4.06 | 1 | d (4.84) | 15 | 1''', 2''', 8, 7 | 76.4 |
| 16 | 4.98 | 1 | dt (9.40, 3.91) | 16 | | 82.0 |
| 17 | | | | | | 181.4 |
| 18 | 5.42 | 1 | d (3.68) | 18 | 10, 4, 13 | 80.3 |
| 19 | 2.34 | 1 | m | 19 | 12 | 31.3 |
| 20 | 1.01 | 3 | d (6.89) | 20 | 5, 19, 21 | 17.0 |
| 21 | 1.13 | 3 | d (6.92) | 21 | 5, 19, 20 | 19.6 |
| 23 | | | | | | 169.7 |
| 1' | 7.67 | 1 | br s | 1' | | 124.5 |
| 2' | 8.09 | 1 | br s | | 1' | 129.7 |
| 3' | 6.62 | 1 | br s | | | 96.1 |
| 4' | | | | | | 133.1 |
| 1" | | | | | | 138.3 |
| 6' | | | | | | 118.0 |
| 7' | | | | | | 162.8 |
| 5' | | | | | | 135.0 |
| 2", 6", 3", 5" | 7.29 | 4 | app d (4.4) | 2", 6", 3", 5" | 4", 1", 7" | 130.3 |
| 4" | 7.22 | 1 | p (4.35) | 4" | | 127.7 |
| 7" | 3.04 | 1 | dd (14.72, 9.41) | 7" | 2", 6", 3", 5", 16 | 36.3 |
| | 2.93 | 1 | dd (14.68, 4.01) | 7" | 2", 6", 3", 5", 16 | |
| 1''' | | | | | | 171.2 |
| 2''' | 2.74 | 2 | m | 2''', 3''' | 1''', 4''', 5''', 6''', 7" | 36.6 |
| 3''' | 2.79 | 2 | m | 2''', 3''' | 1''', 4''', 3''' | 16.6 |
| 4''' | | | | | | 83.7 |
| 5''' | | | | | | 74.9 |
| 6''' | | | | | | 66.4 |
| 7''' | | | | | | 75.8 |
| 8''' | | | | | | 123.2 |
| 9''', 13''' | 7.46 | 2 | m | 9''', 13''' | 9''', 13''', 8''', 7''' | 133.4 |
| 10''', 12''' | 7.35 | 2 | m | 10''', 12''' | 8''', 10''', 12''' | 129.5 |
| 11''' | 7.38 | 1 | t t (7.4, 1.3) | 11''' | | 130.1 |
| (NH) | 1.89 | 1 | br s | | | |

### Supplementary Figures

### A

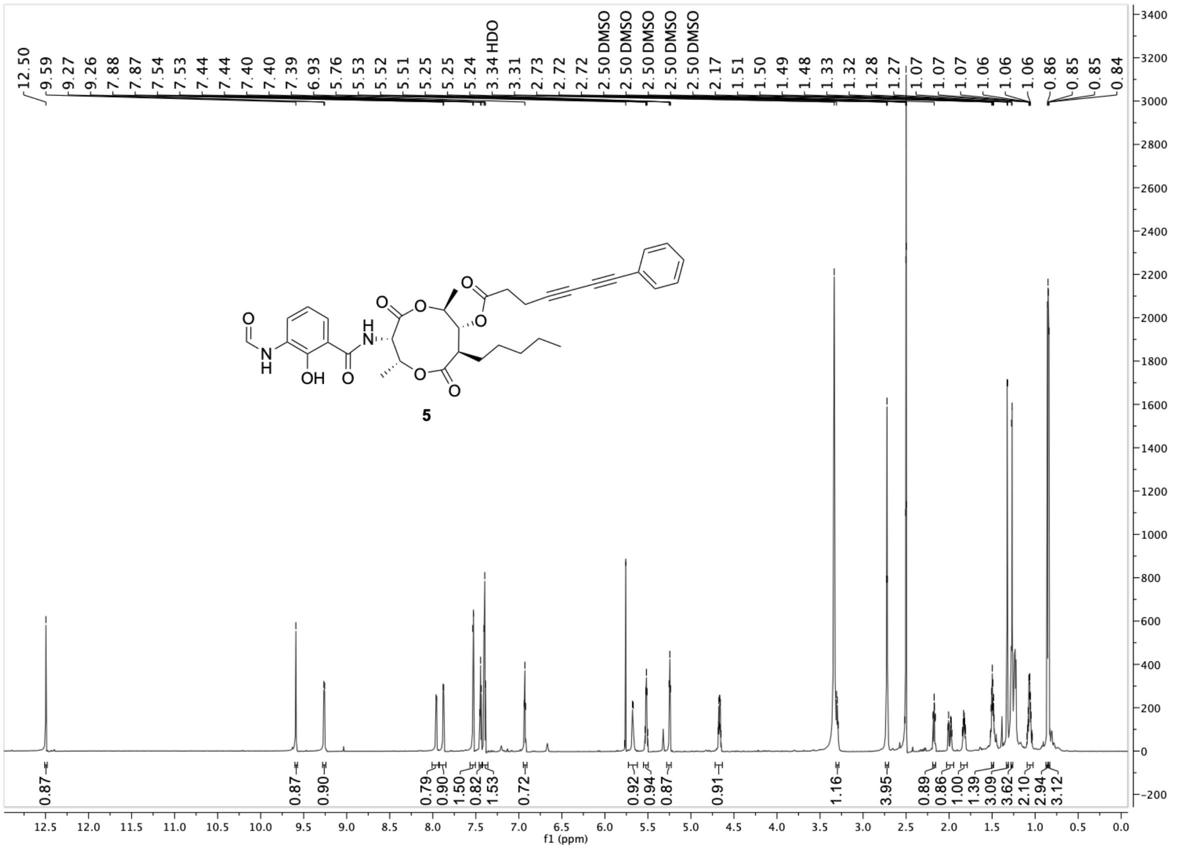

**Figure S1:** NMR, MS, and UV-vis characterization of PhDY-Ant (**5**). A) $^1H$ NMR spectrum (DMSO-$d_6$, 900 MHz) of **5**.

B

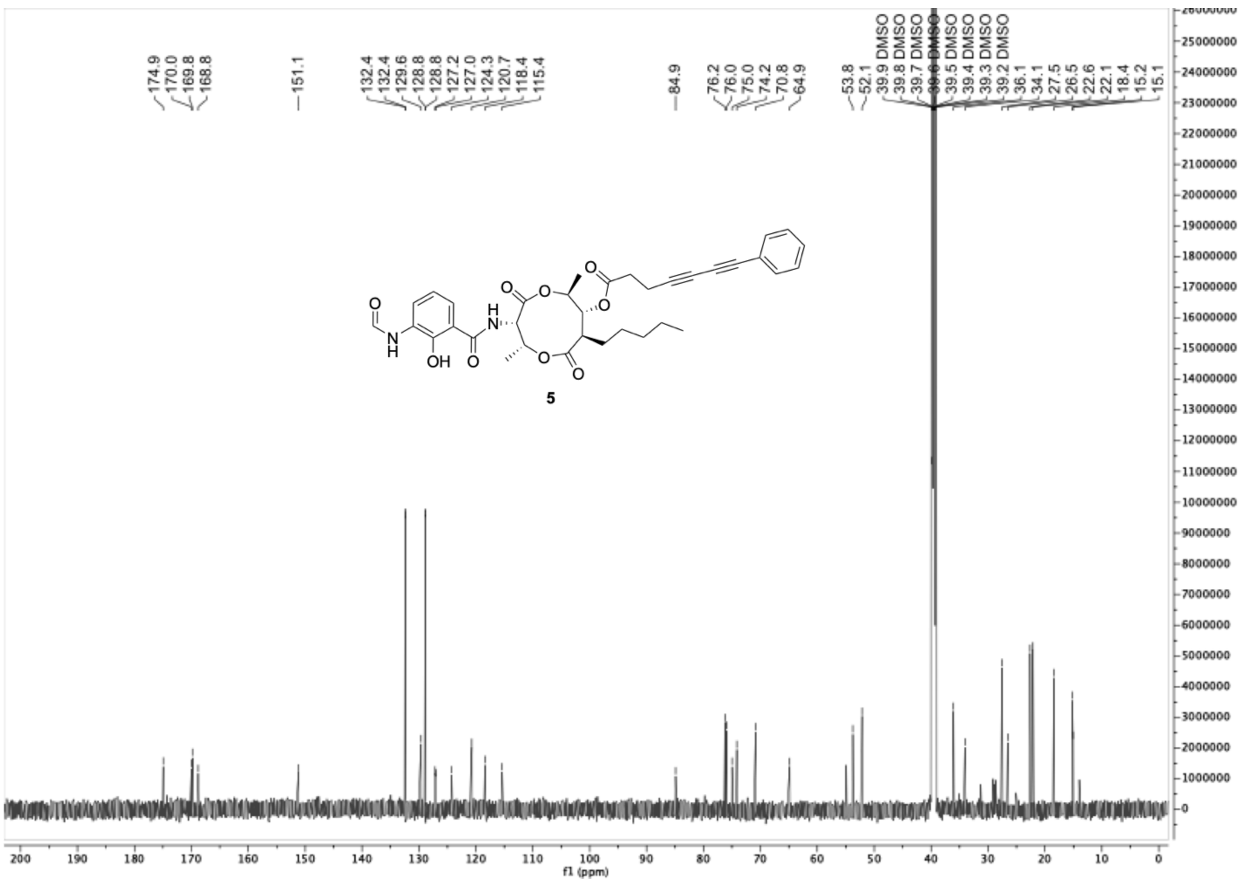

**Figure S1:** B) $^{13}C$ NMR spectrum (DMSO-$d_6$, 226 MHz) of **5**.

**C**

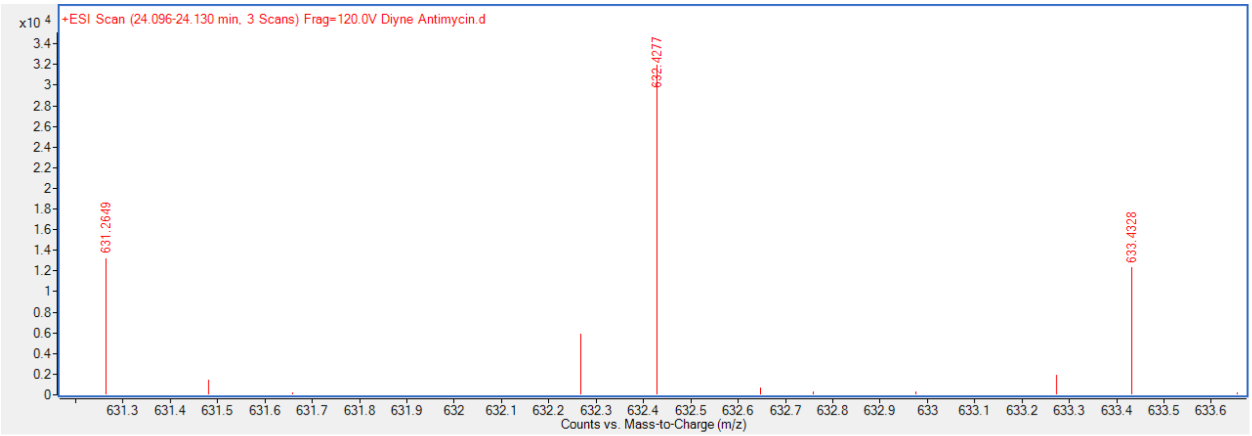

**Figure S1:** C) Mass spectrum of **5**. Calcd for $C_{49}H_{55}N_2O_{13}$ $[M+H]^+$ *m/z* 631.2650, found 631.2649.

**D**

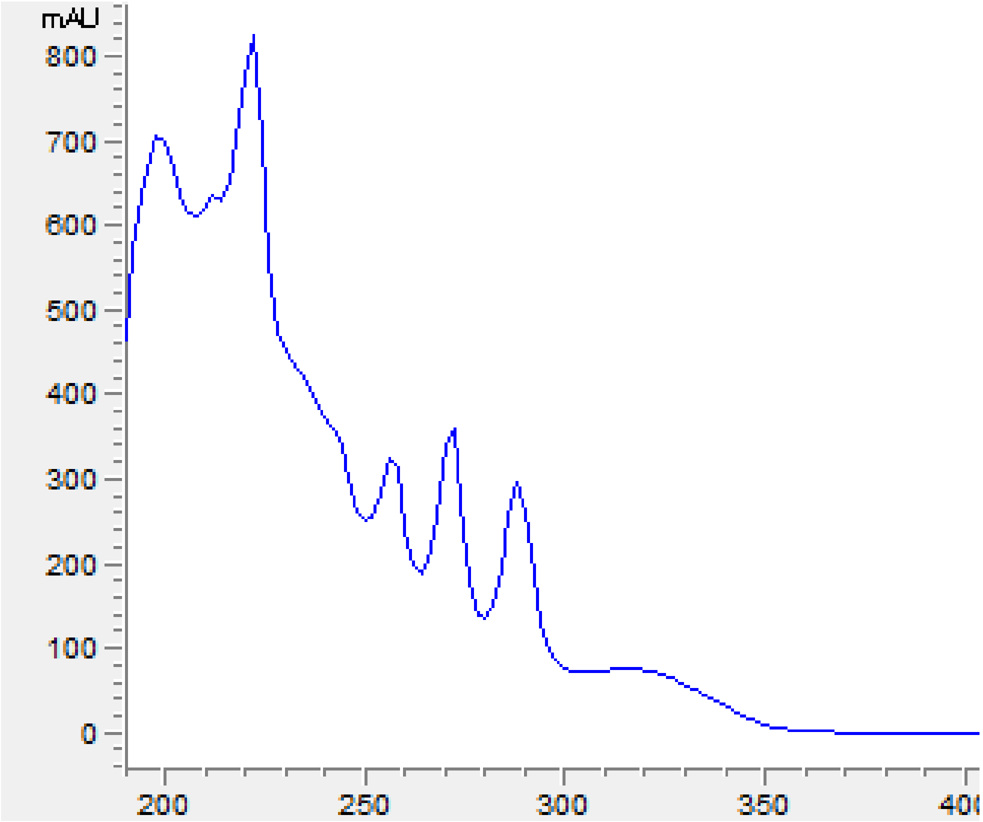

**Figure S1:** D) UV-vis spectrum of **5**.

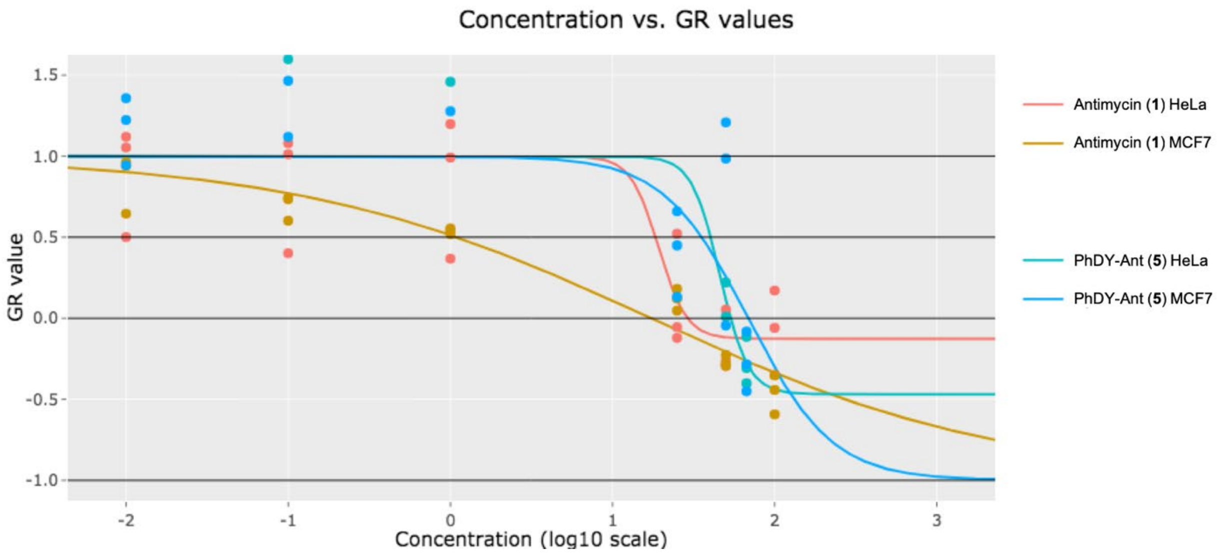

**Figure S2.** PhDY-Ant (**5**) and antimycin (**1**) dose response curves. Antimycin (100 μM to 0.010 μM, red and gold), and PhDY-Ant (**5**) (67 μM to 0.010 μM, teal and blue lines) dose response curves in HeLa (red and teal lines) and MCF-7 (gold and blue lines) cells show the PhDY Raman tag minimally perturbs activity. Concentration of compounds is in μM (log scale). Data were collected in triplet and growth response curves were generated as described above. The curve shown is representative of at least three biological replicates.

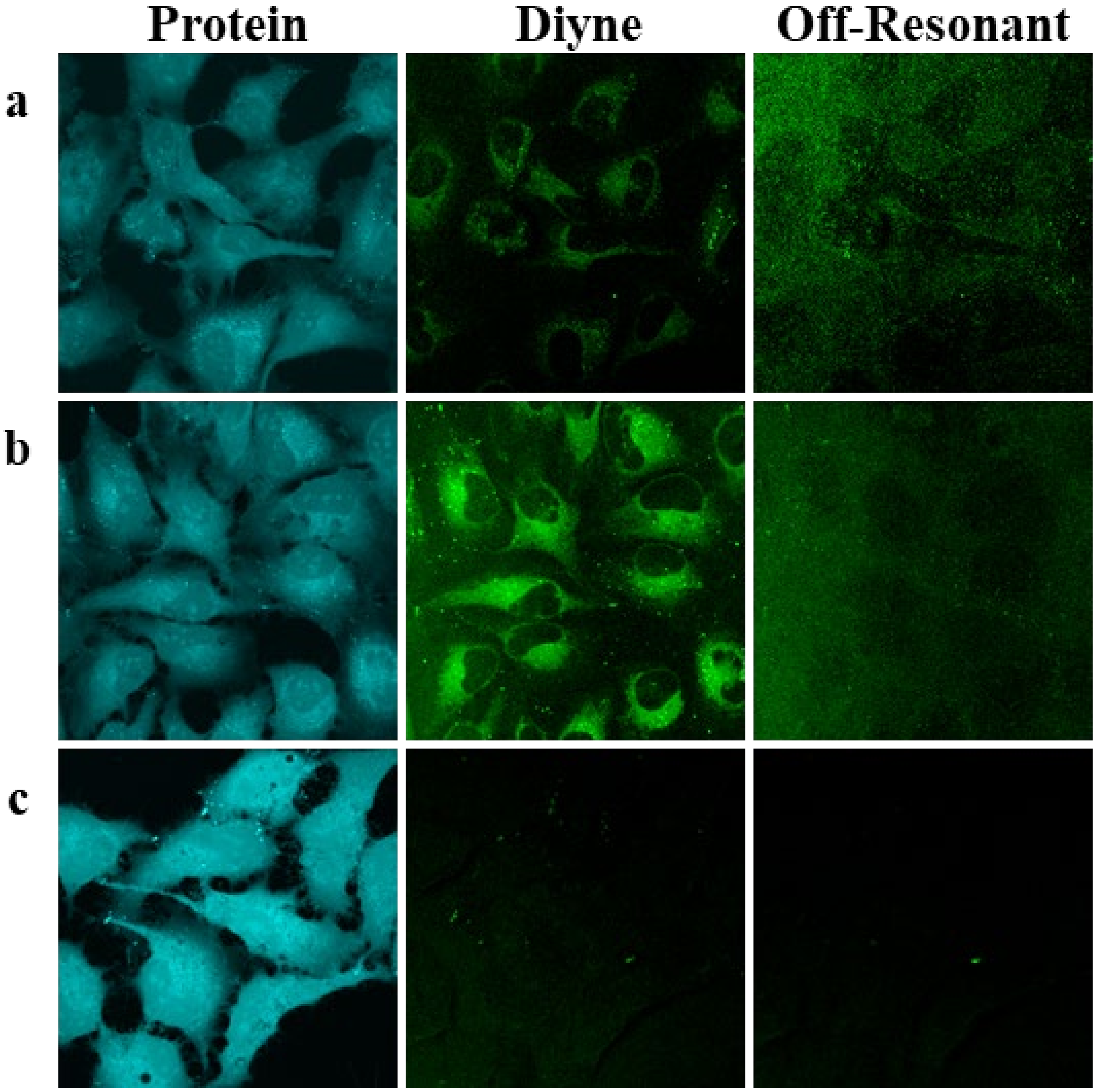

**Figure S3.** Intracellular PhDY-Ant (**5**) and phenyl-diyne tag imaging at various concentrations in HeLa cells. Cells were imaged 2 h post PhDY-Ant addition at 10 μM (a) and 50 μM (b). Phenyl-diyne tag (50 μM) was imaged as a negative control (c). SRS images were acquired by tuning the frequency difference between the pump and Stokes lasers to be resonant with protein ($CH_3$, 2940 $cm^{-1}$), diyne (2251 $cm^{-1}$), off-resonance (2000 $cm^{-1}$).

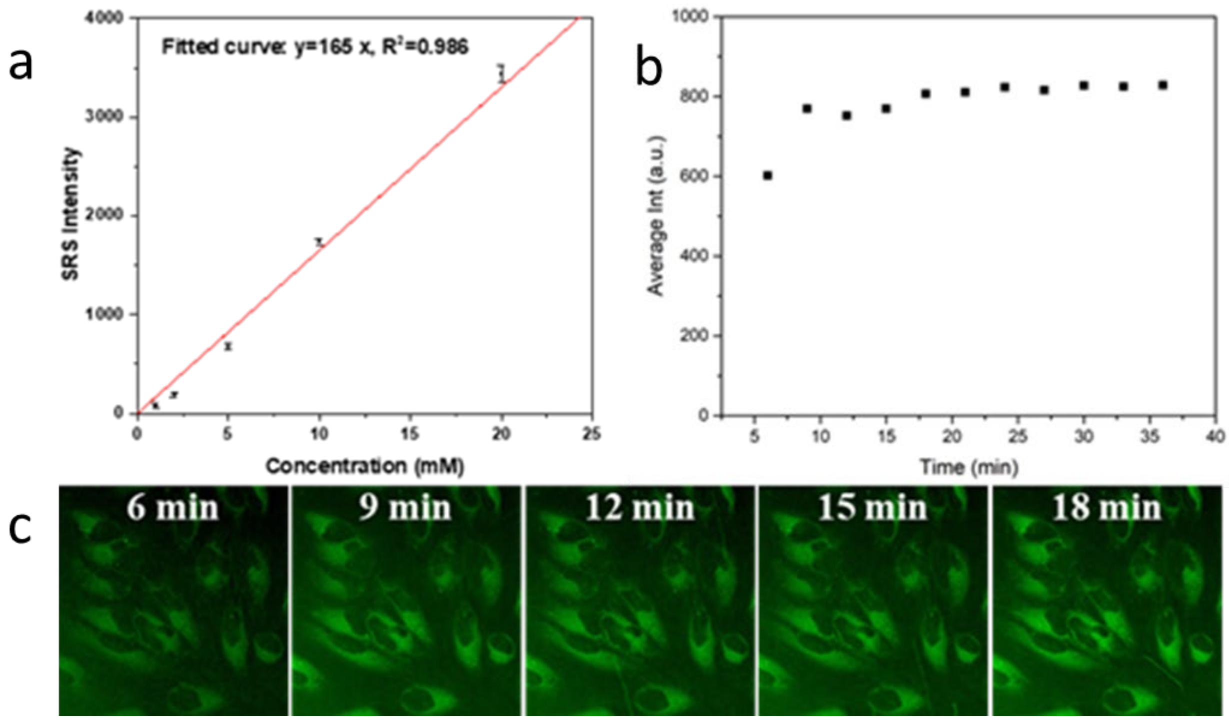

**Figure S4.** Time course of PhDY-Ant (**5**) uptake in HeLa cells. a) Calibration curve showing relationship of PhDY-Ant concentration with signal intensity. b) Average signal intensity over time. c) SRS images used to analyze the time course experiment. HeLa cells incubated with 50 μM PhDY-Ant (**5**).

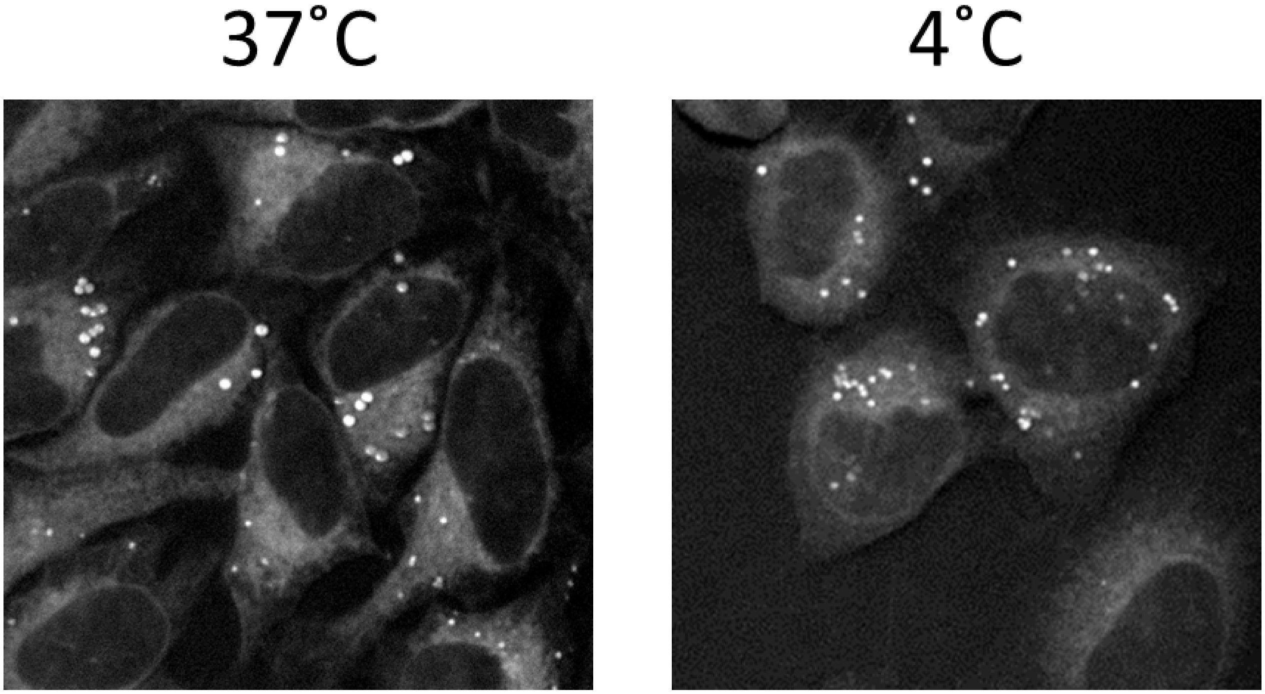

**Figure S5.** Temperature dependent uptake of PhDY-Ant (**5**). SRS images showing uptake of 50 µM **5** during 2h incubation at 4˚C and 37 ˚C. **5** was still absorbed into HeLa cells at 4˚C, indicating that the uptake mechanism is energy-independent.

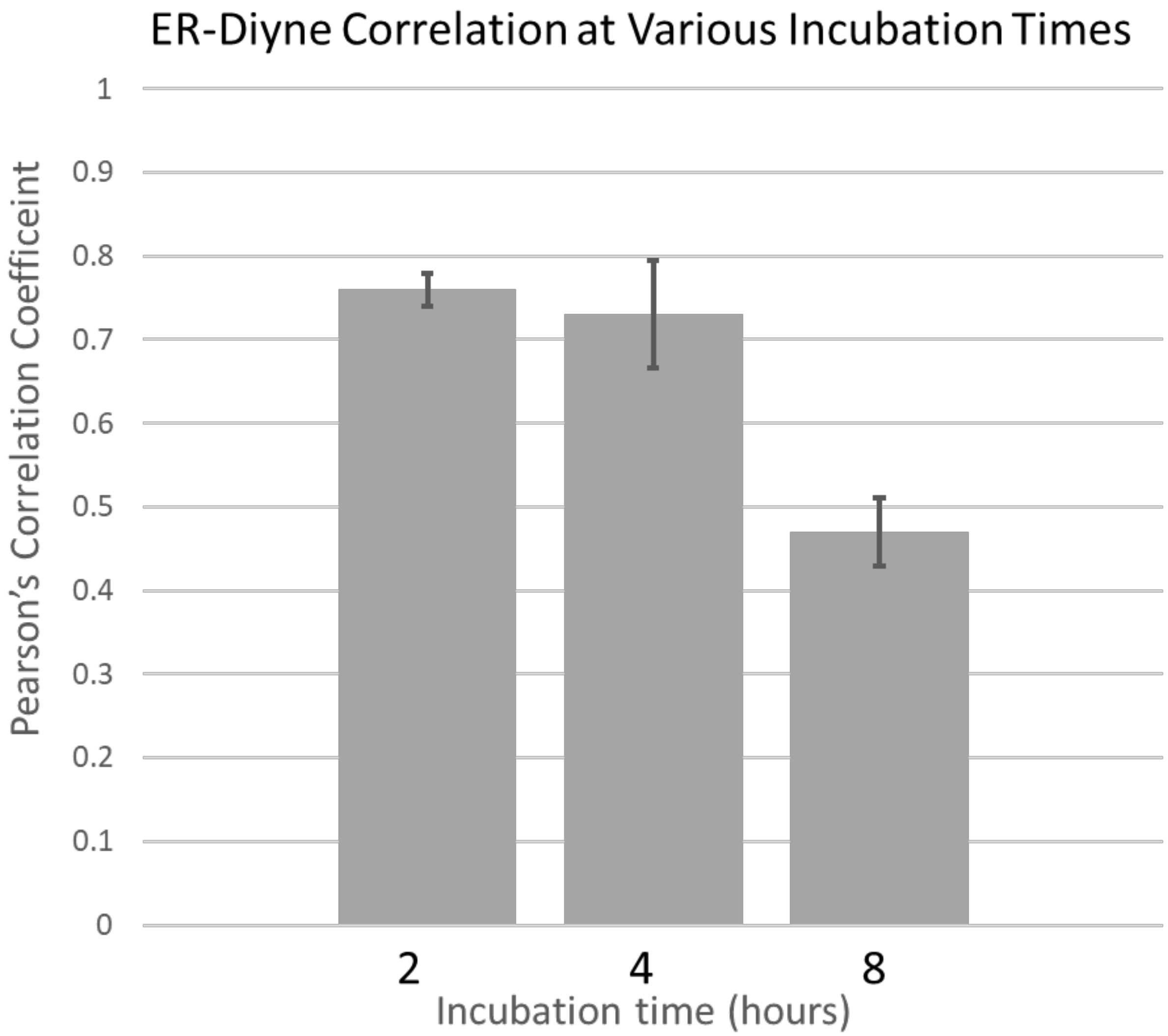

**Figure S6.** Time-dependent colocalization of PhDY-Ant (**5**) in HeLa cells. Pearson's correlation coefficients were calculated using the coloc2 plugin for imageJ in triplicate.

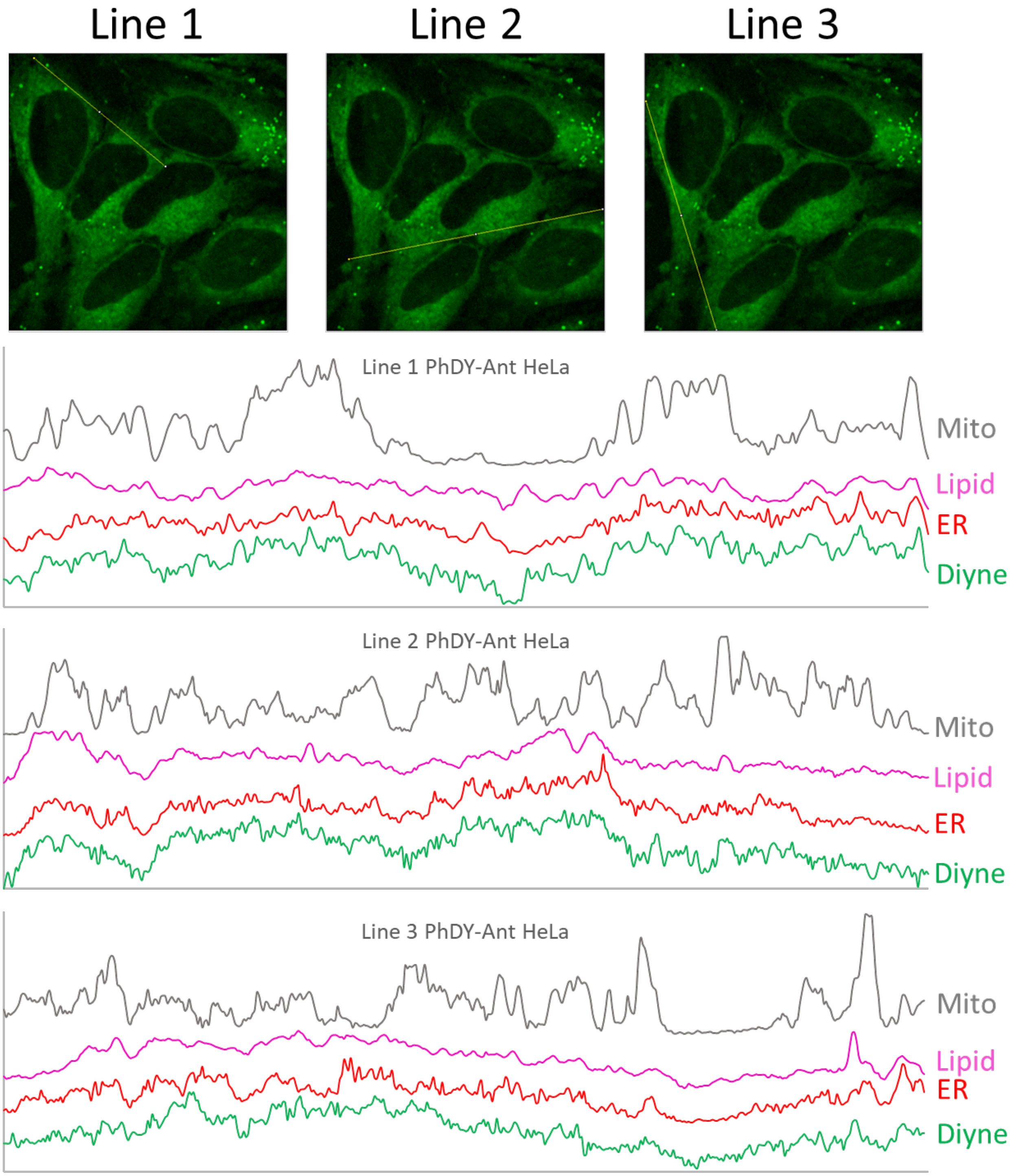

**Figure S7.** Line plot analysis of PhDY-Ant (**5**) in HeLa cells. Each line plot shows poor correlation between Mito-tracker and the diyne channel. Each line plot shows good correlation between the lipid channel and the diyne channel. The ER-tracker and diyne channels are very well-correlated, including some features that are excluded on the lipid channel.

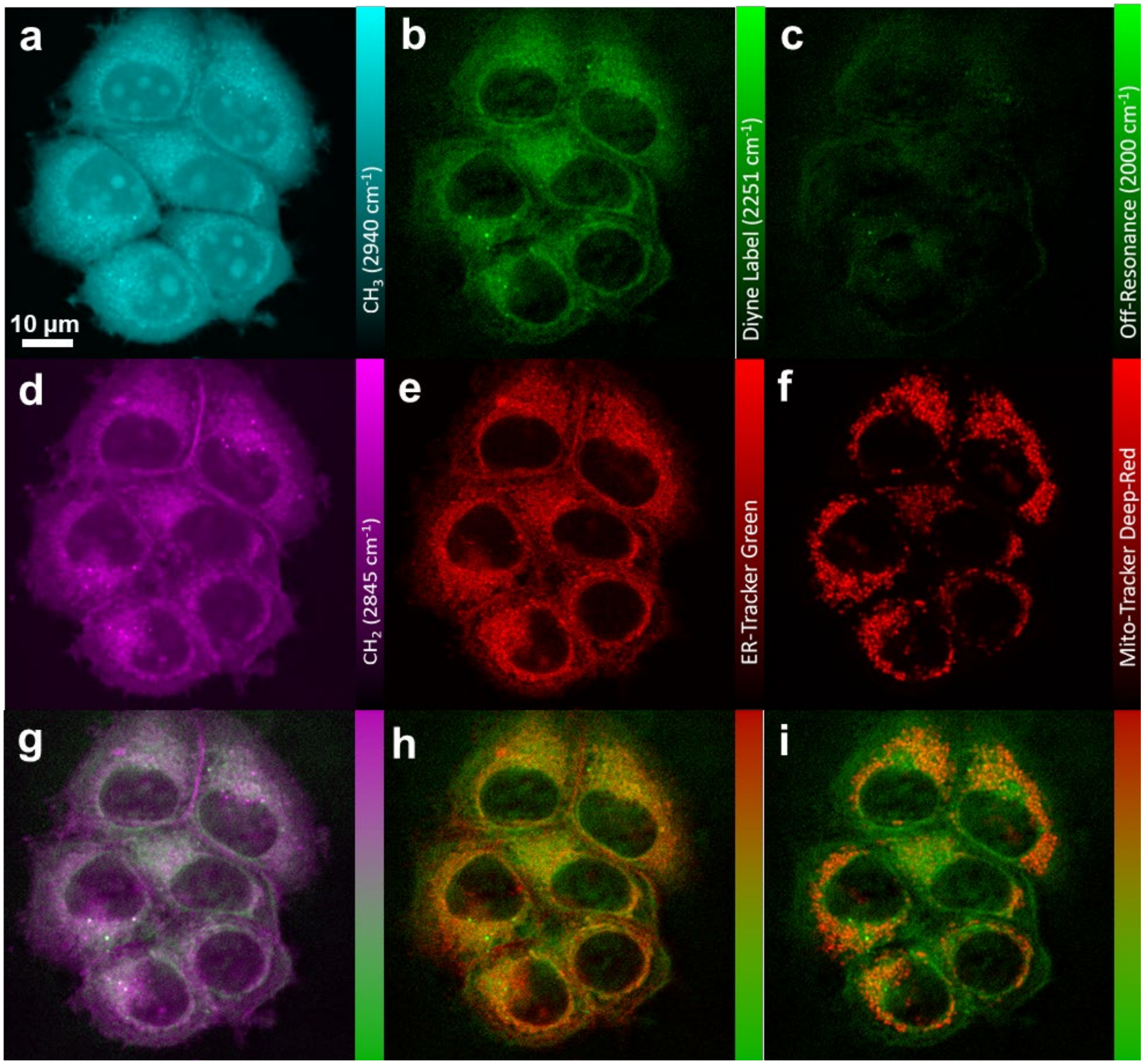

**Figure S8.** SRS and fluorescence imaging of PhDY-Ant (**5**) in MCF-7 cells. **a)** $CH_3$ channel at 2940 $cm^{-1}$ representing proteins. **b)** Diyne label at 2251 $cm^{-1}$. **c)** Off-resonance channel at 2000 $cm^{-1}$. **d)** $CH_2$ channel at 2845 $cm^{-1}$, representing lipids. **e)** Confocal fluorescence imaging of ER-Tracker excited at 488 nm. **f)** Confocal fluorescence imaging of Mito-Tracker excited at 635 nm. **g)** Overlay image of d) lipids and b) diyne label. **h)** Overlay image of e) ER-Tracker and b) diyne label. **i)** Overlay image of f) Mito-Tracker and b) diyne label.

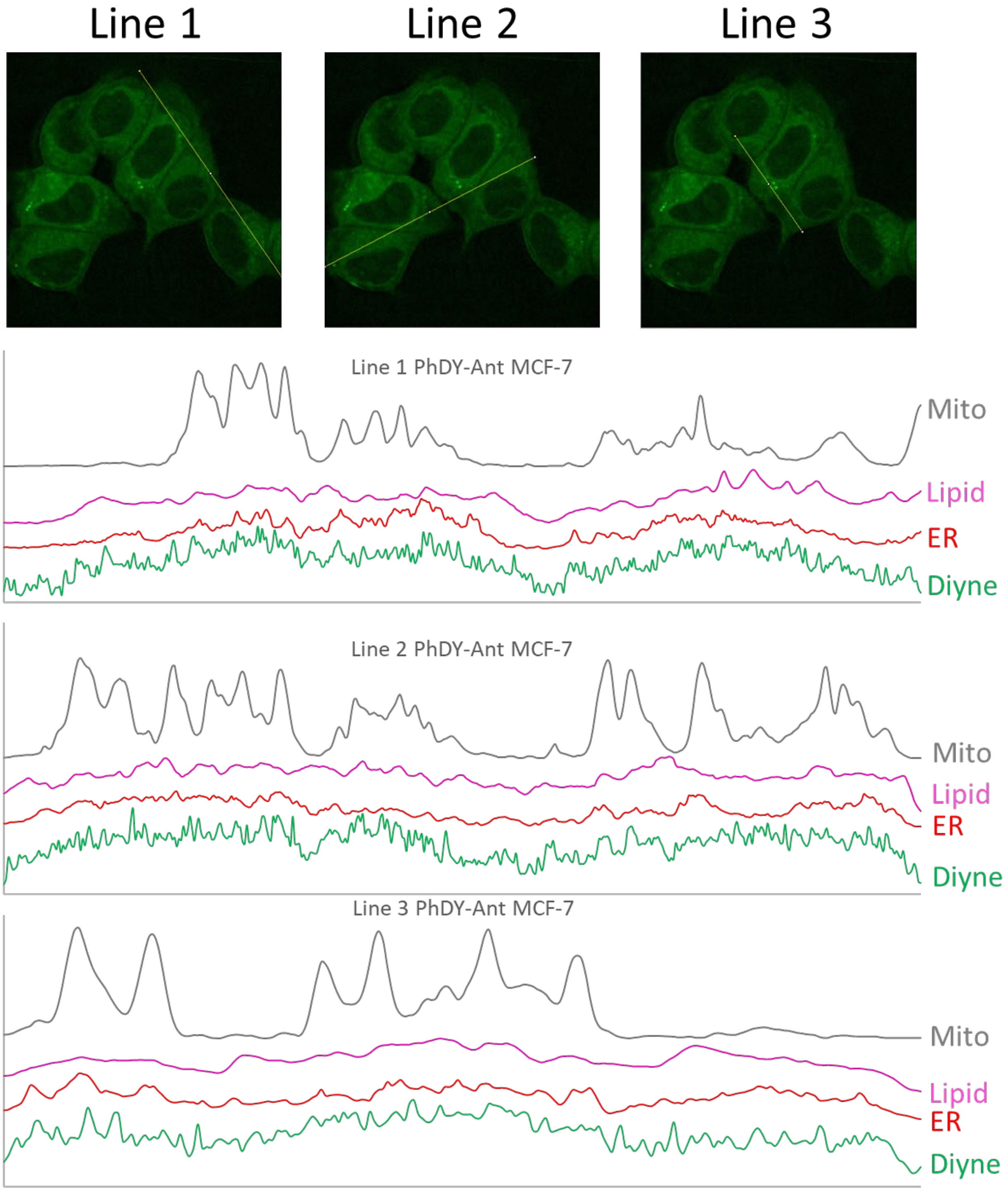

**Figure S9.** Line plot analysis of PhDY-Ant (**5**) in MCF-7 cells. Each line plot shows poor correlation between Mito-tracker and the diyne channel. The features of the diyne channel can be explained by a combination of the lipid and ER-tracker cannels.

**A**

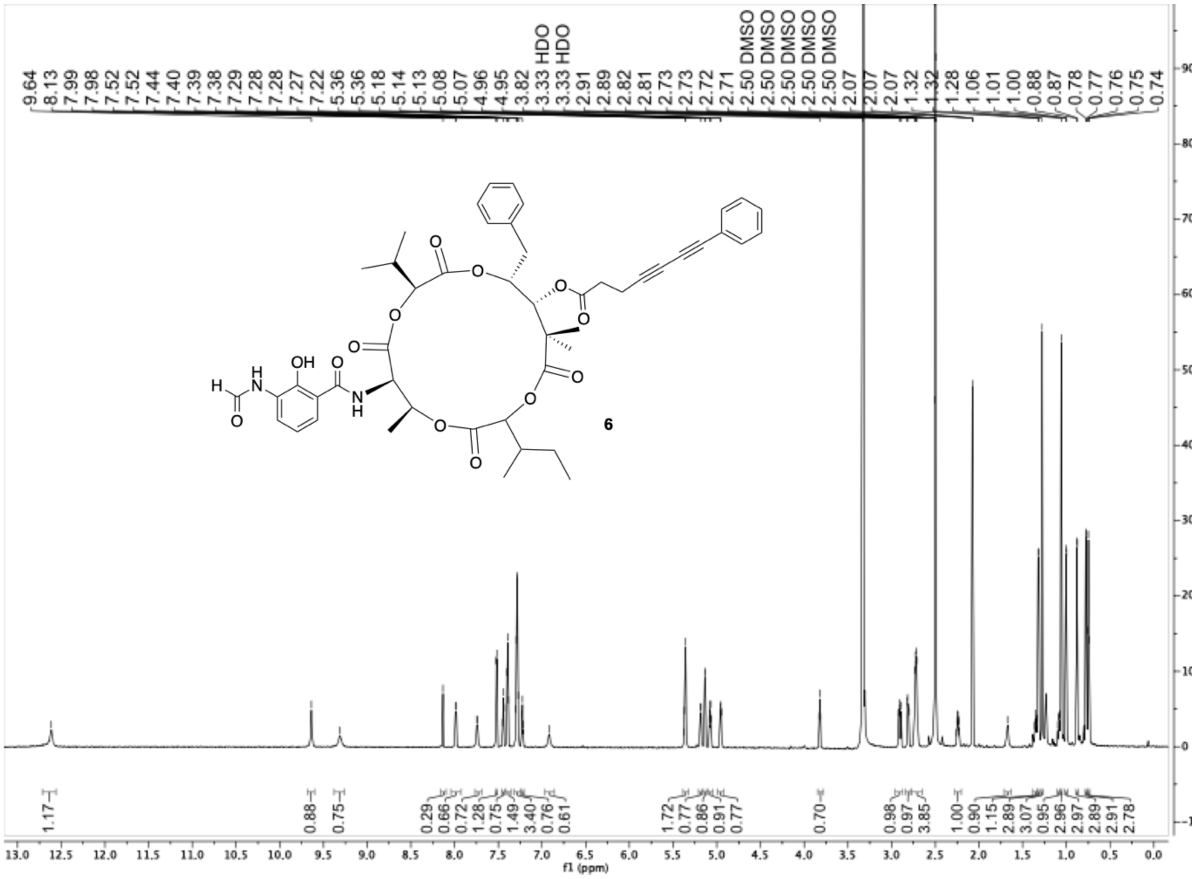

**Figure S10.** NMR, MS, and UV-vis characterization of PhDY-NeoA (**6**). A) $^{1}H$ NMR spectrum (DMSO-$d_6$, 900 MHz) of **6**.

**B**

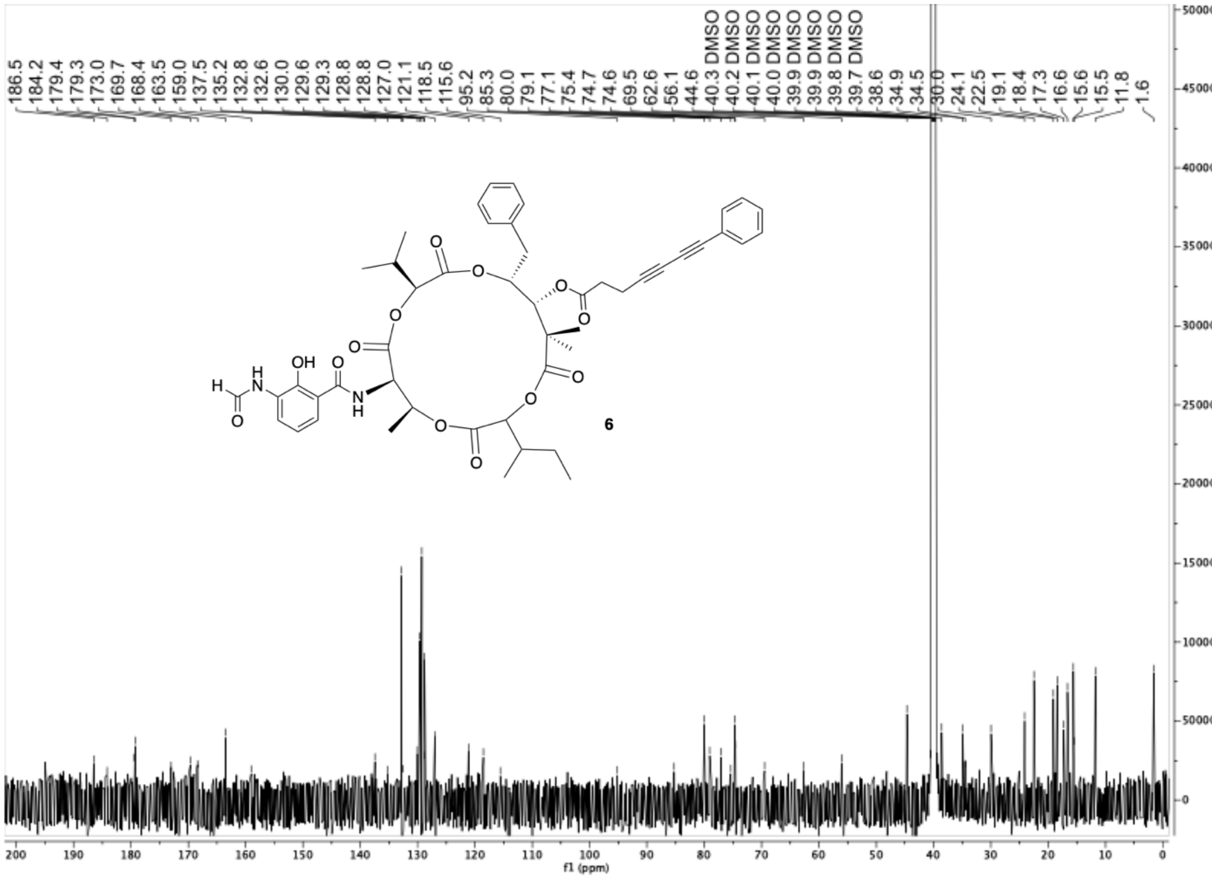

**Figure S10.** B) $^{13}$C NMR spectrum (DMSO-$d_6$, 900 MHz) of **6**.

**C**

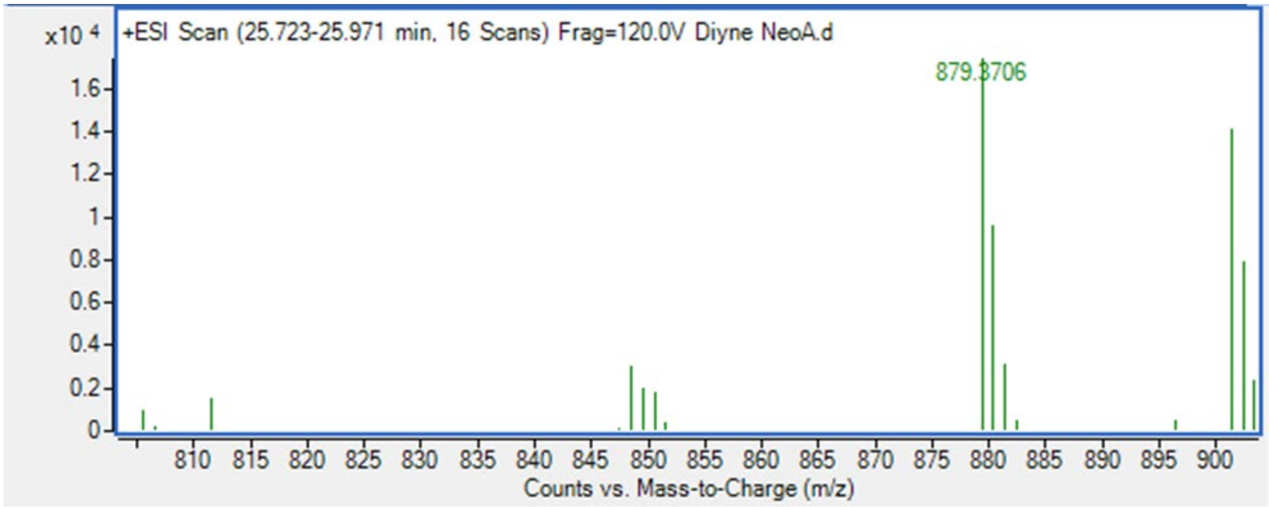

**Figure S10.** C) Mass spectrum of **6**. Calcd for $C_{49}H_{55}N_2O_{13}$ $[M+H]^+$ *m/z* 879.3699, found 879.3706.

**D**

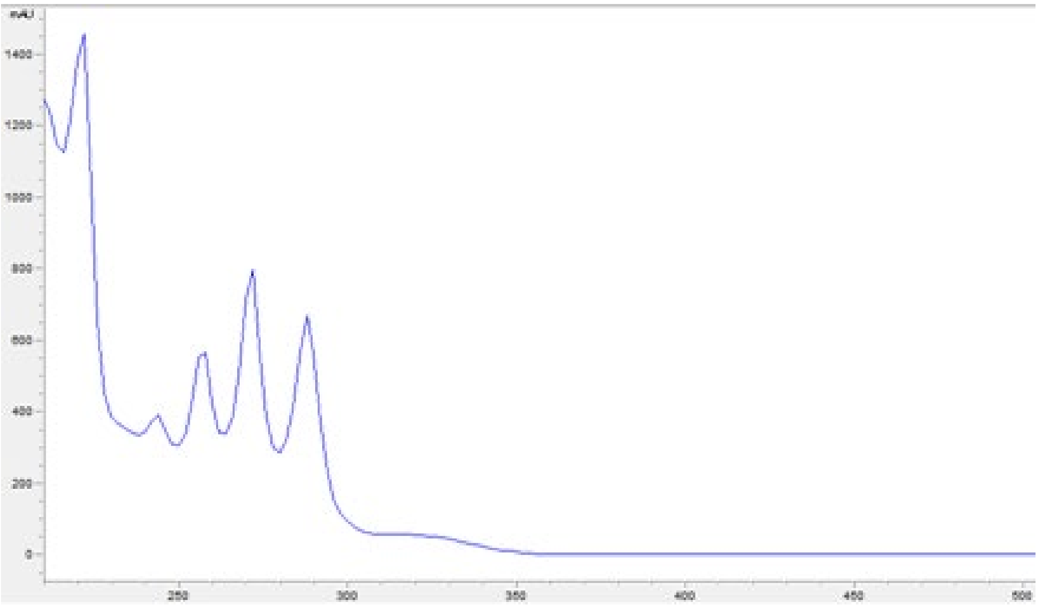

**Figure S10.** D) UV-vis spectrum of **6**.

A

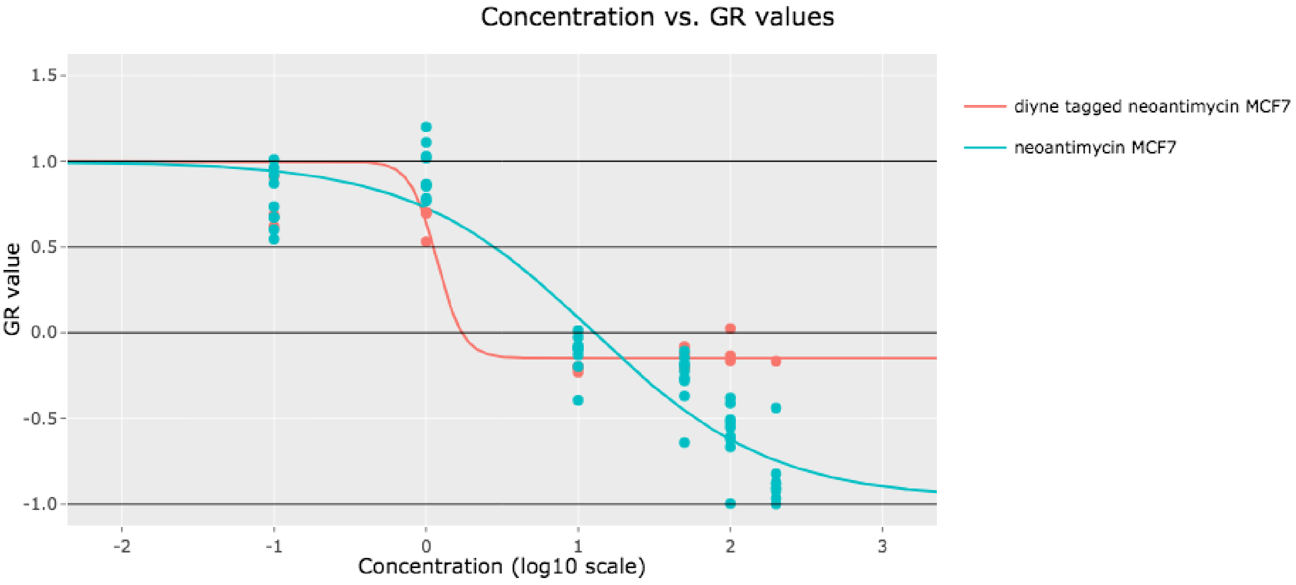

B

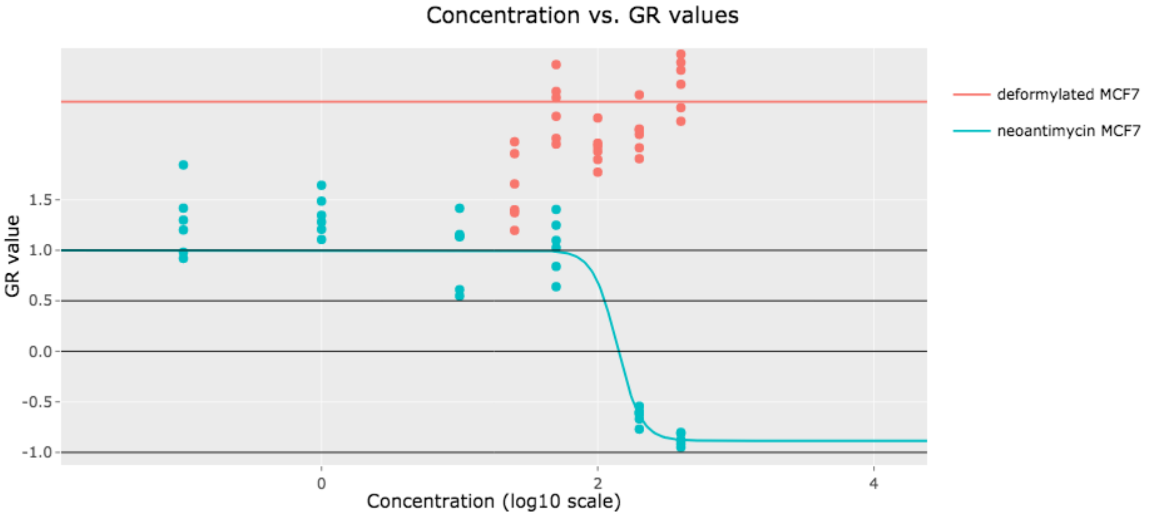

**Figure S11.** Dose response curve for various neoantimycins in MCF-7 cells. Concentration of compounds is in μM (log scale). Data were collected in triplet to sextuplet and growth response curves were generated as described in methods. The curves shown are representative of three individual biological replicates. A) Neoantimycin (**2**) and PhDY-NeoA (**6**) (200 μM to 0.10 μM), dose response curves in MCF-7 cells show the Raman-active tag at C-11 is minimally perturbing. The plateau at higher concentrations of PhDY-NeoA is most likely due to solubility limits of the compound. B) Neoantimycin (**2**) (1.0 mM to 0.10 μM), and deformylated neoantimycin (**7**) (400 μM to 25 μM) dose response curves in MCF-7 cells demonstrate the importance of the *N*-formyl group.

A

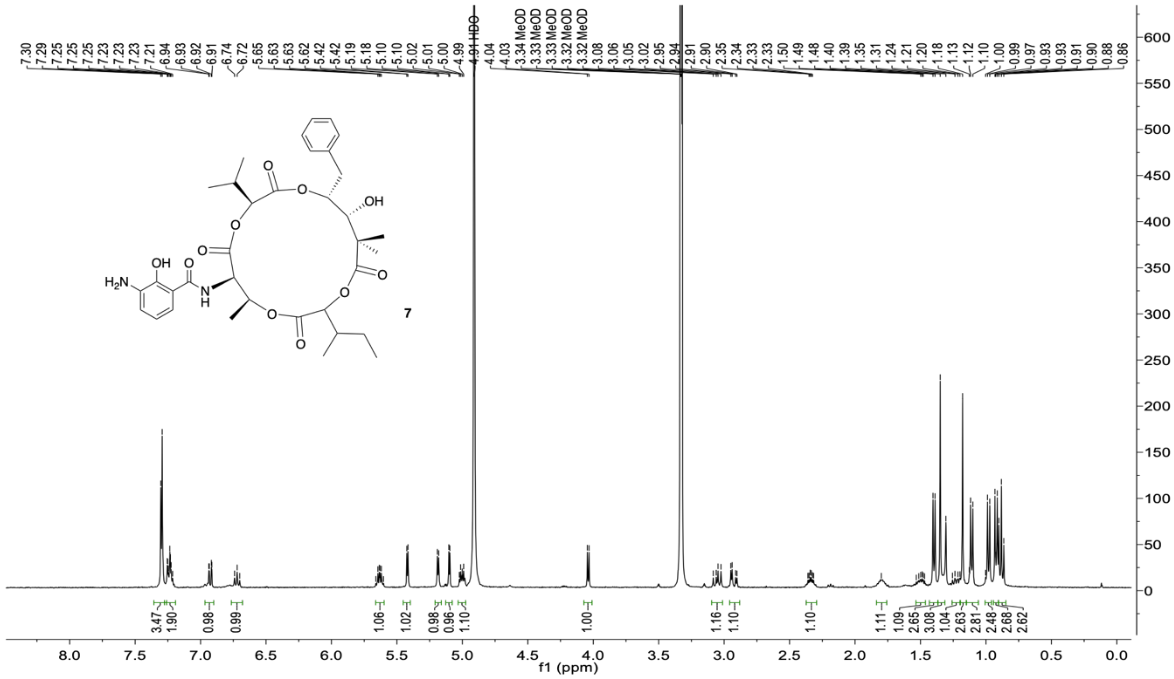

**Figure S12.** NMR, MS, and UV-vis characterization of deformylated neoantimycin (**7**). A) $^{1}H$ NMR spectrum ($CD_3OD$, 900 MHz) of **7**.

B

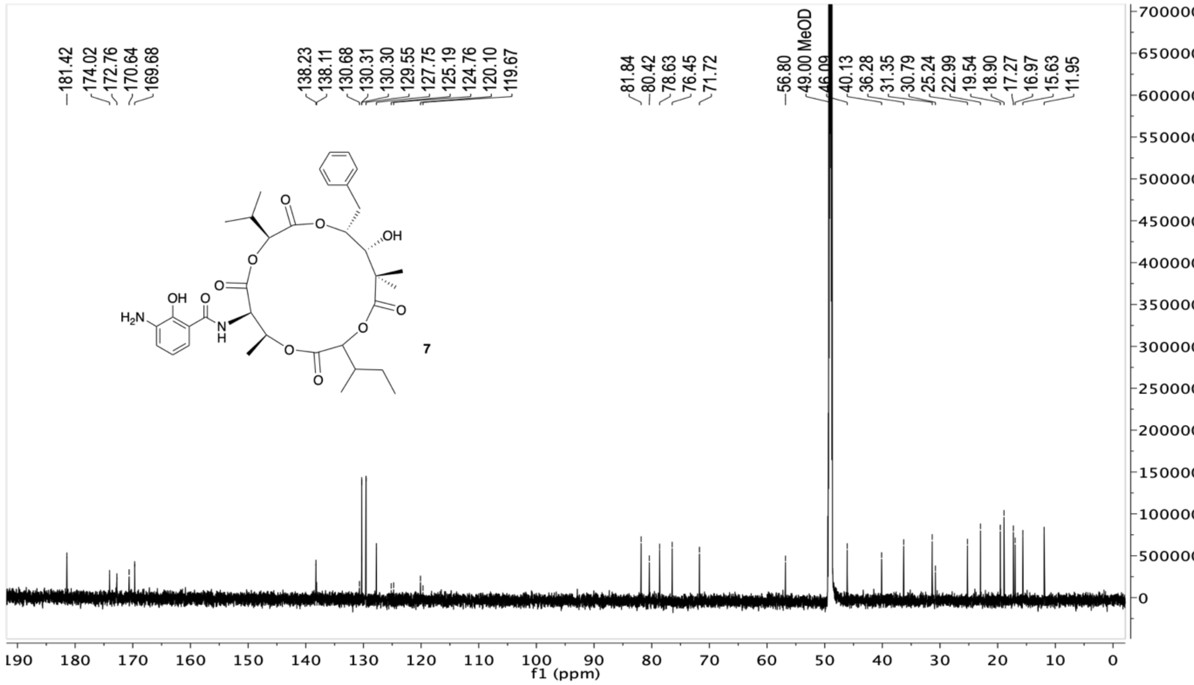

**Figure S12.** B) $^{13}C$ NMR spectrum ($CD_3OD$, 226 MHz) of **7**.

**C**

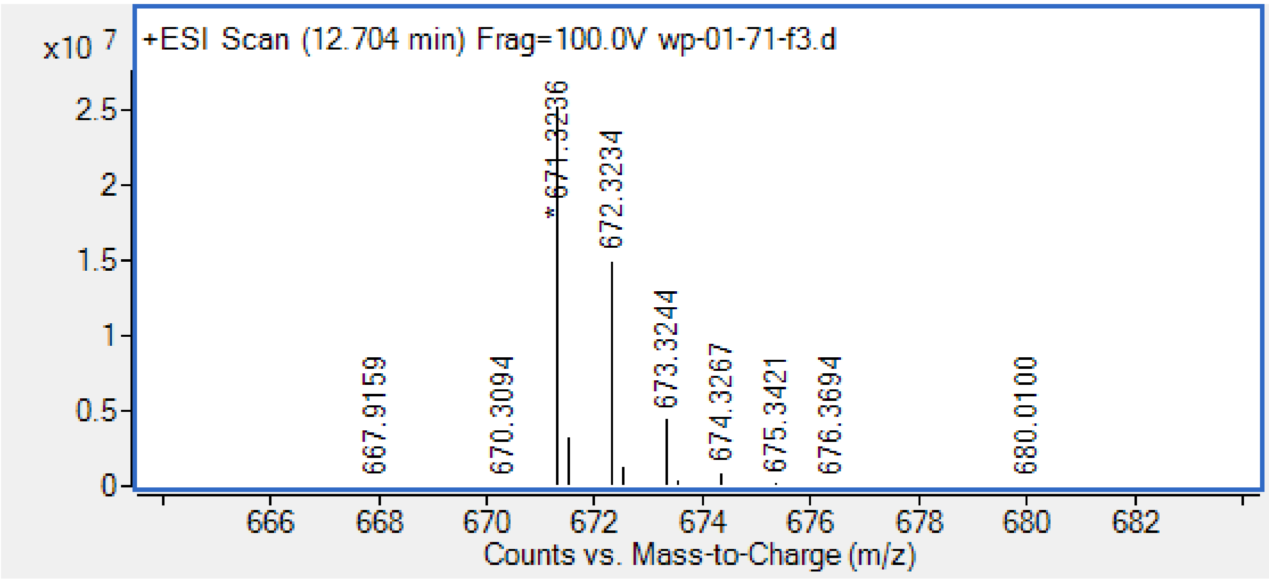

**Figure S12.** C) Mass spectrum of compound **7**. Calcd for $C_{35}H_{47}N_2O_{11}$ $[M+H]^+$ *m/z* 671.3174, found 671.3236.

**D**

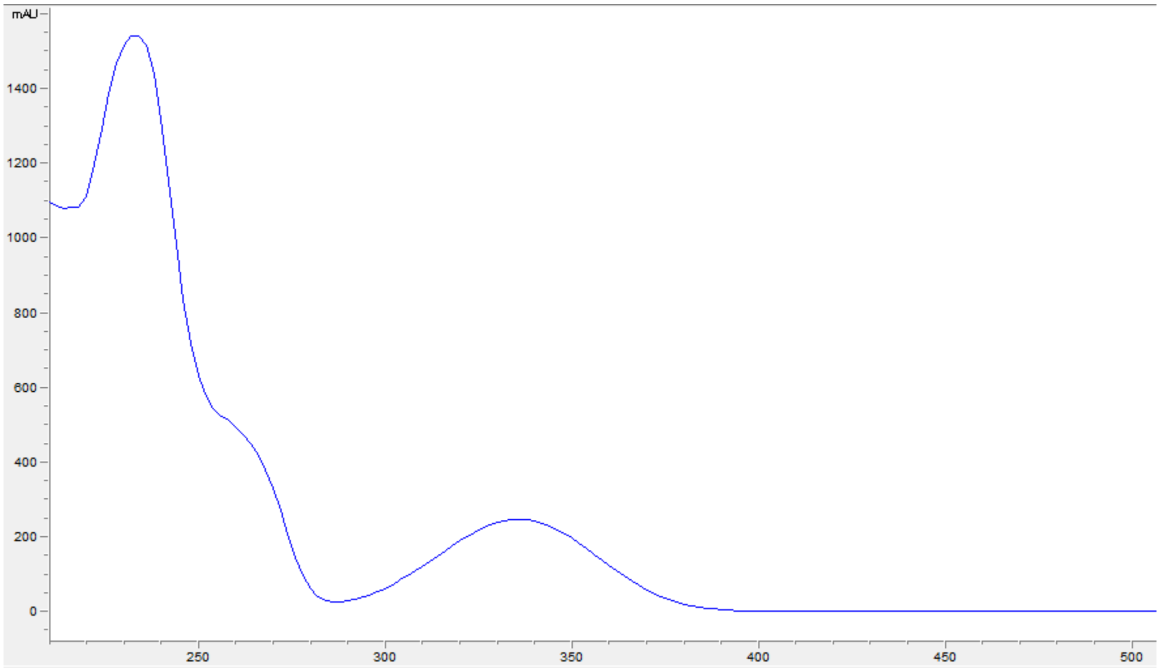

**Figure S12.** D) UV-vis spectrum of **7**.

**A**

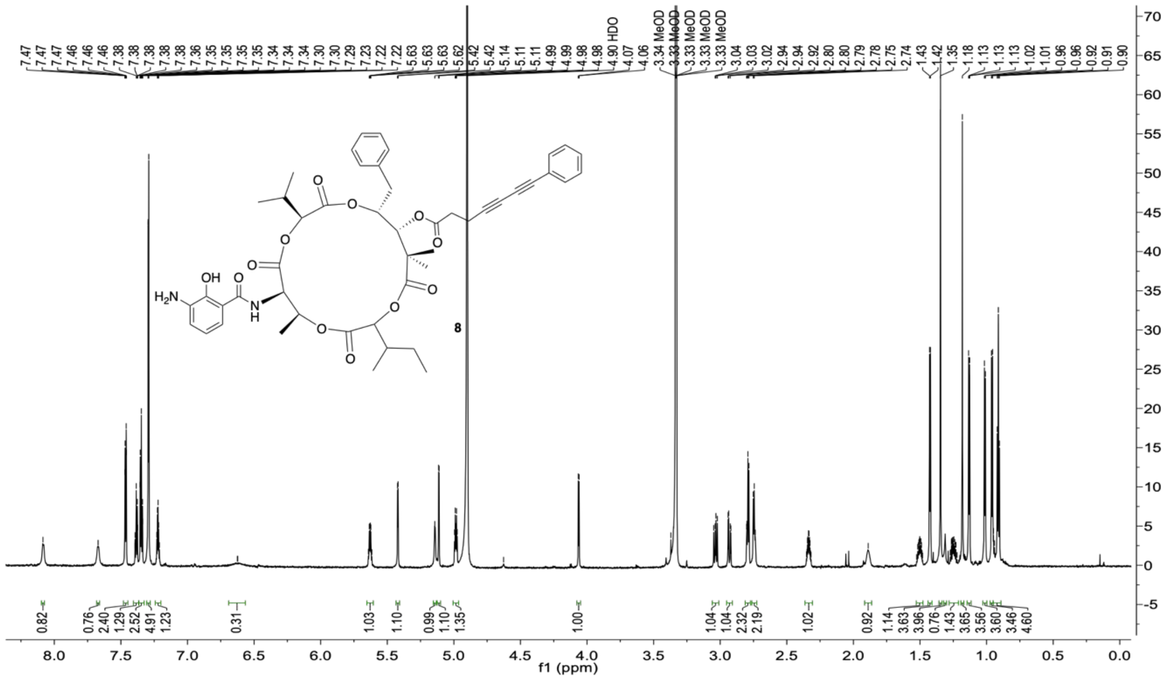

**Figure S13.** NMR, MS and UV-vis characterization of deformylated PhDY-NeoA (**8**). A) [1]H NMR $CD_3OD$, 900 MHz) of **8**.

**B**

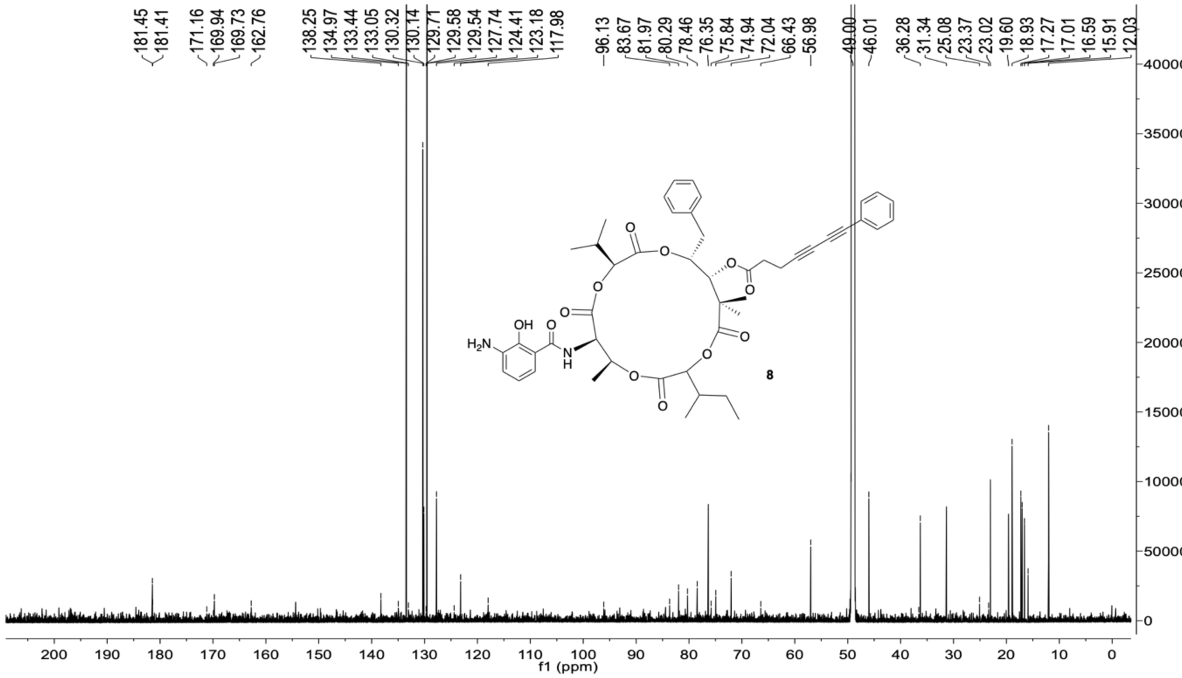

**Figure S13.** B) [13]C NMR ($CD_3OD$, 226 MHz) of **8**.

C

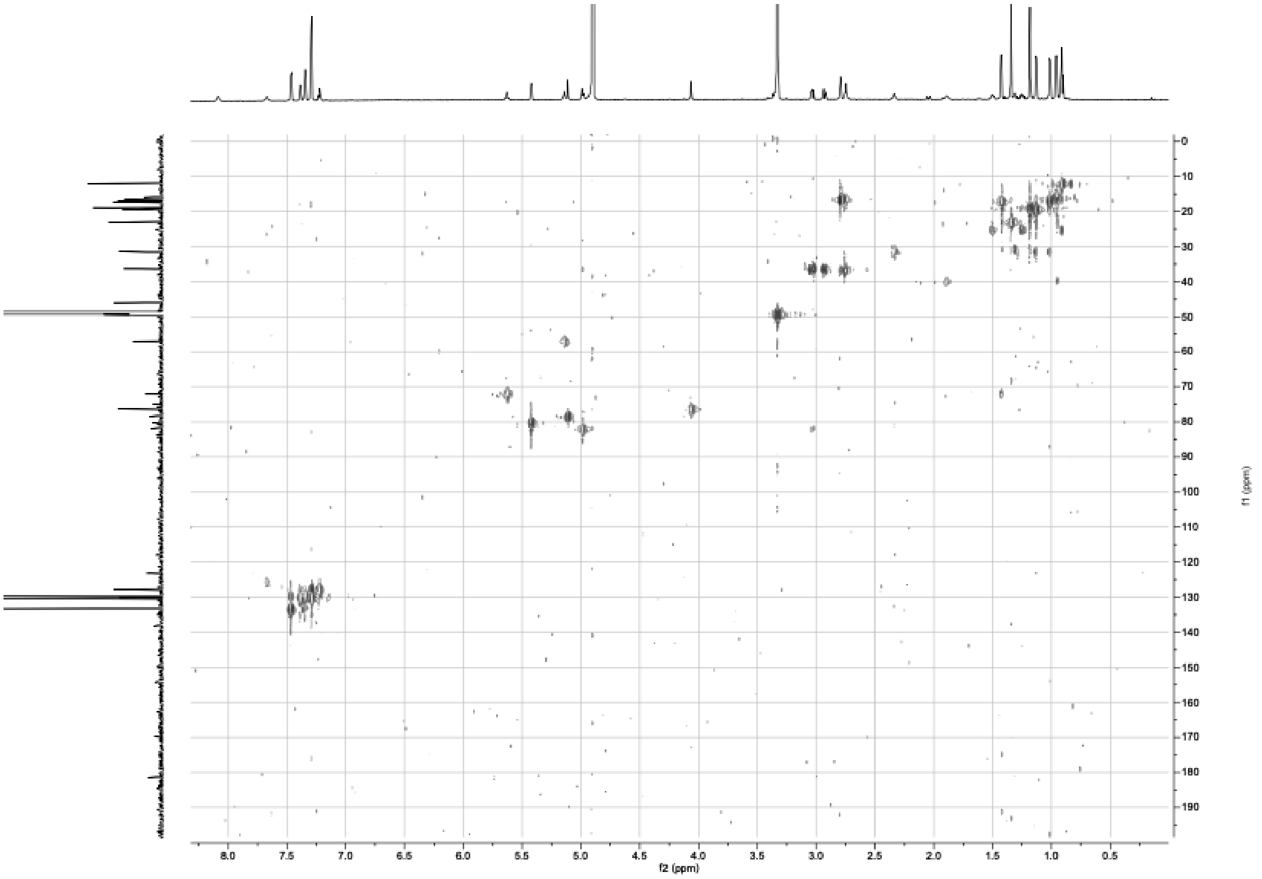

**Figure S13.** C) HSQC spectrum ($CD_3OD$, 900 MHz) of **8.**

D

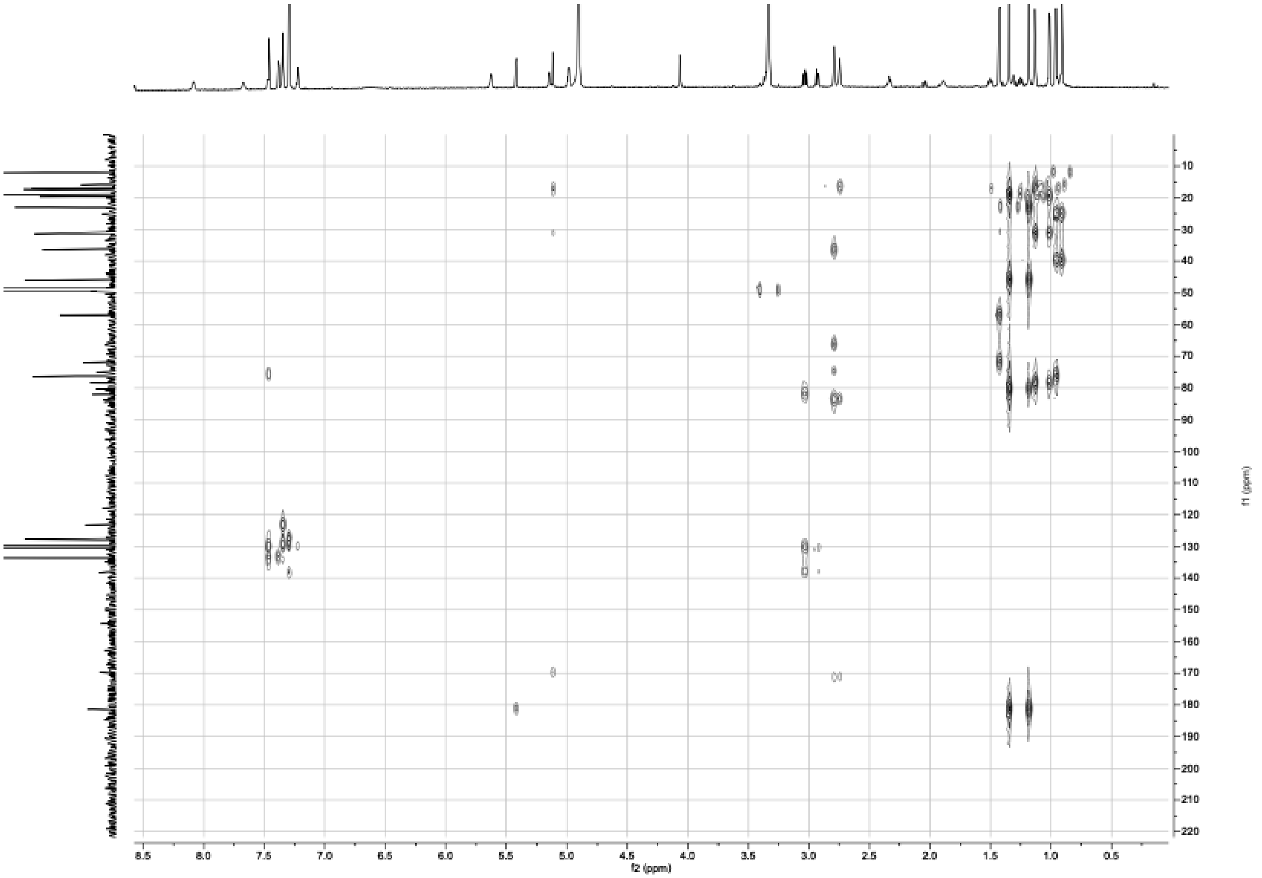

**Figure S13.** D) HMBC spectrum ($CD_3OD$, 900 MHz) of **8.**

E

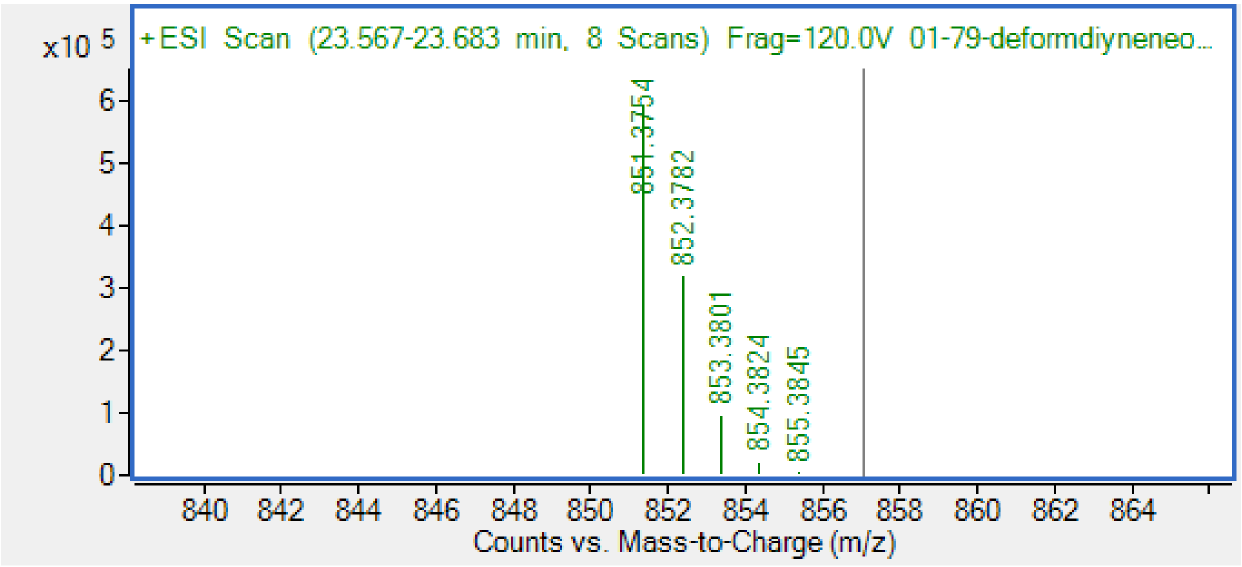

**Figure S13.** E) Mass spectrum of **8**. Calcd for $C_{48}H_{55}N_2O_{12}$ $[M+H]^+$ *m/z* 851.3750, found 851.3754.

F

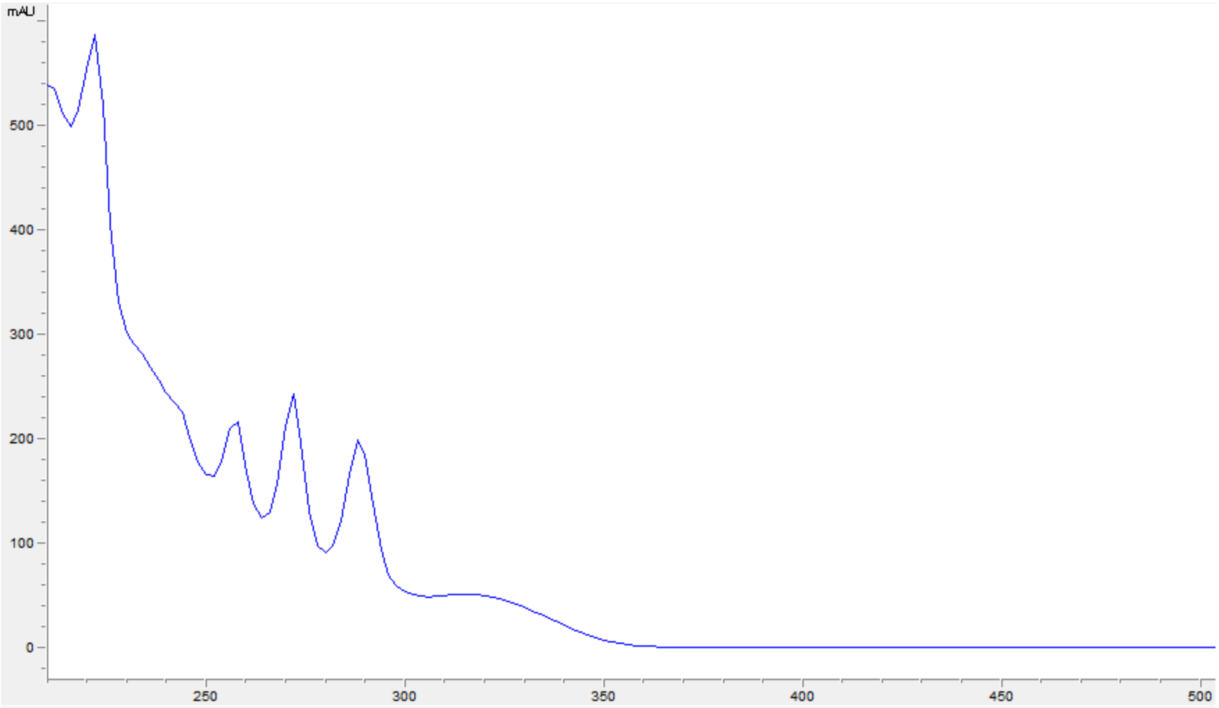

**Figure S13.** F) UV-vis spectrum of compound (**8**).

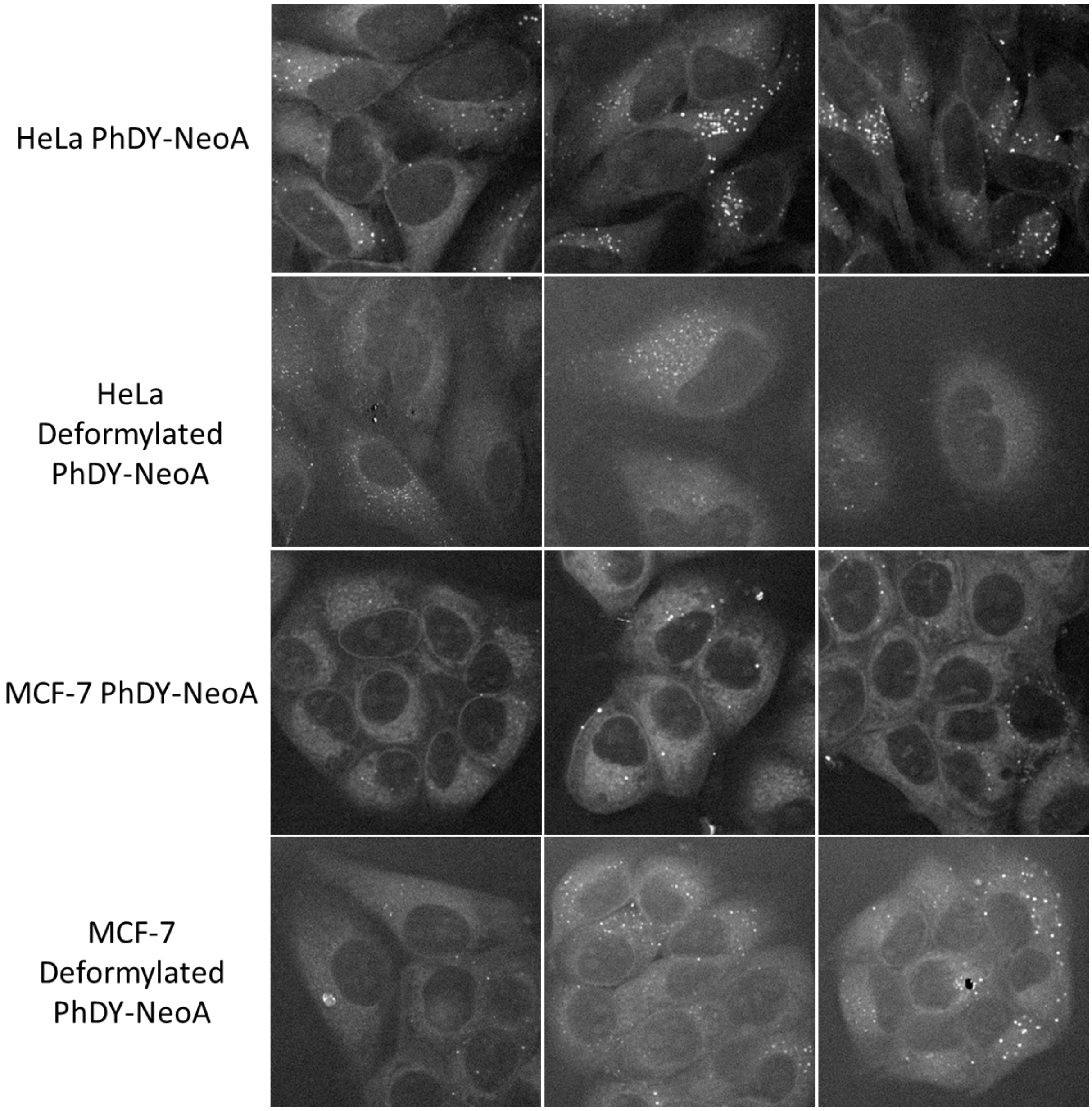

**Figure S14.** Imaging of differential uptake of PhDY-NeoA (**6**) and deformylated PhDY-NeoA (**8**) in HeLa and MCF-7 cells.

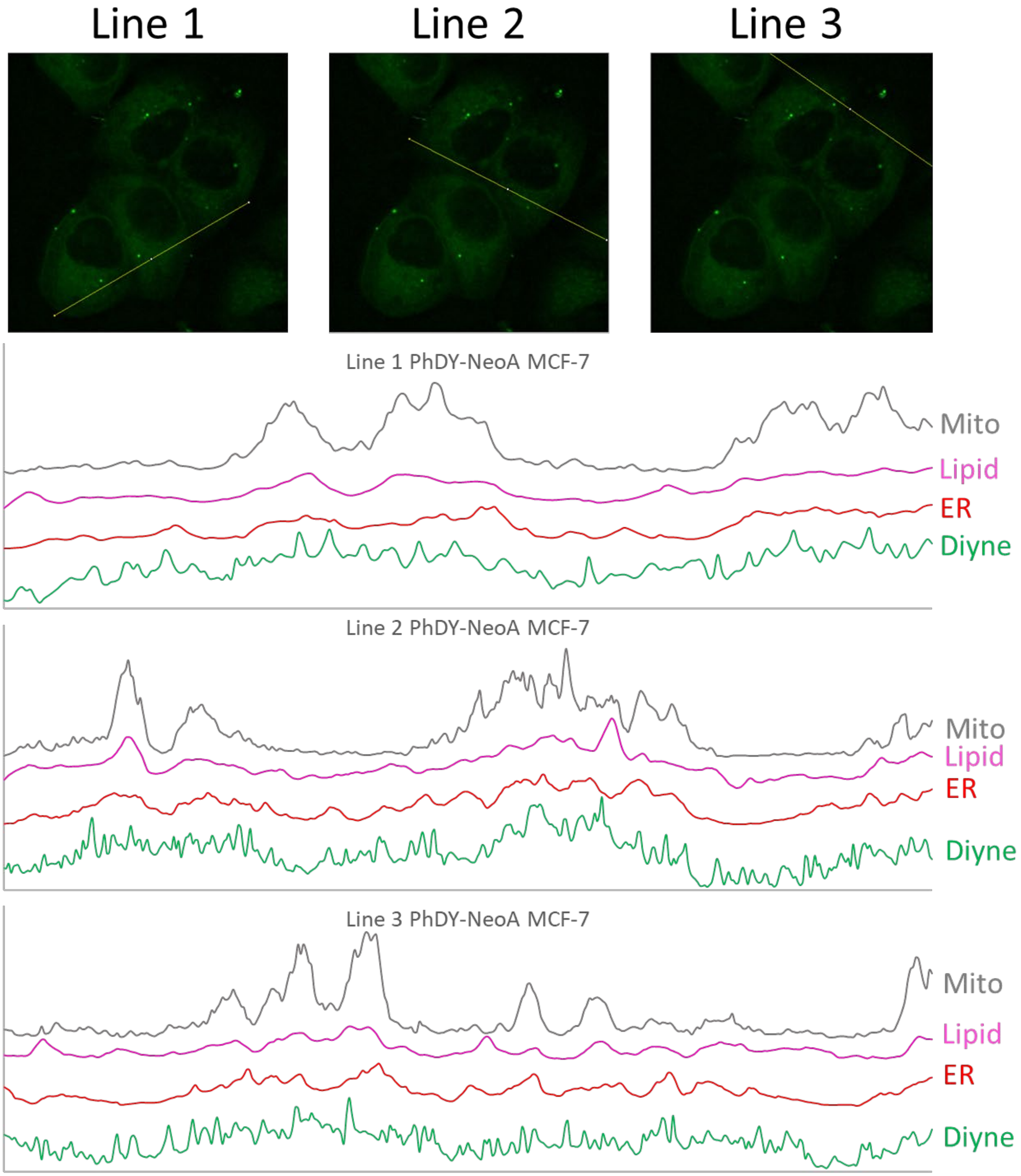

**Figure S15.** Line plot analysis of PhDY-NeoA (**6**) in MCF-7 cells. Each line plot shows poor correlation between Mito-tracker and the diyne channel. Each line plot shows good correlation between the lipid channel and the diyne channel. The ER-tracker and diyne channel are less well-correlated than the diyne-lipid pair.
